# BayeSMART: Bayesian Clustering of Multi-sample Spatially Resolved Transcriptomics Data

**DOI:** 10.1101/2024.08.30.610571

**Authors:** Yanghong Guo, Bencong Zhu, Chen Tang, Ruichen Rong, Ying Ma, Guanghua Xiao, Lin Xu, Qiwei Li

**Affiliations:** The University of Texas at Dallas; Chinese University of Hong Kong; The University of Texas Southwestern Medical Center; Brown University

**Author notes:** **Corresponding authors:** Guanghua Xiao, Quantitative Biomedical Research Center, Peter O’Donnell Jr. School of Public Health, The University of Texas Southwestern Medical Center, Dallas, Texas, USA., Lin Xu, Quantitative Biomedical Research Center, Peter O’Donnell Jr. School of Public Health, The University of Texas Southwestern Medical Center, Dallas, Texas, USA., Qiwei Li, Department of Mathematical Sciences, The University of Texas at Dallas, Richardson, Texas,USA.

**Keywords:** AI-reconstructed histology image, Markov random field, multi-sample analysis, spatial clustering, spatial domain identification

## Abstract

The field of spatially resolved transcriptomics (SRT) has greatly advanced our understanding of cellular microenvironments by integrating spatial information with molecular data collected from multiple tissue sections or individuals. However, methods for multi-sample spatial clustering are lacking, and existing methods primarily rely on molecular information alone. This paper introduces BayeSMART, a Bayesian statistical method designed to identify spatial domains across multiple samples. BayeSMART leverages artificial intelligence (AI)-reconstructed single-cell level information from the paired histology images of multi-sample SRT datasets while simultaneously considering the spatial context of gene expression. The AI integration enables BayeSMART to effectively interpret the spatial domains. We conducted case studies using four datasets from various tissue types and SRT platforms and compared BayeSMART with alternative multi-sample spatial clustering approaches and a number of state-of-the-art methods for single-sample SRT analysis, demonstrating that it surpasses existing methods in terms of clustering accuracy, interpretability, and computational efficiency. BayeSMART offers new insights into the spatial organization of cells in multi-sample SRT data.

## INTRODUCTION

The development of spatially resolved transcriptomics (SRT) technologies provides a new way of measuring biological insights at the cellular level while retaining the spatial layout of tissues [1-3]. One category of SRT methods builds upon next-generation sequencing (NGS)-based SRT techniques using spatial barcoding, including spatial transcriptomics (ST) [4], 10x Visium, Slide-seq [5], Slide-seqV2 [6], and HDST [7]. Each barcoded area (i.e., spot) typically contains multiple cells, while Slide-seq, Slide-seqV2, and HDST further enhance resolution by measuring transcripts at a near-cellular or sub-cellular level. Another category is imaging-based technologies, characterized by methods such as fluorescence *in situ* hybridization (FISH) or sequencing, including seqFISH [8], MERFISH [9], and STARmap [10]. These methods often measure genes at single-cell resolution. The advancements in SRT have significantly enhanced the knowledge of cell interactions, spatial heterogeneity in the disease state, and complex multicellular biological systems [11-13].

Spatial domain identification (also known as spatial clustering) is one of the fundamental challenges in SRT data [14, 15]. Among the existing clustering methods on single-sample SRT data, the Seurat package [16, 17] utilizes the high-throughput gene expression without considering the spatial information. The hidden-Markov random field model (HMRF) [18] and BayesSpace [19] integrate spatial information under the Bayesian frameworks but fail to utilize the paired histology images. To incorporate histological information into spatial domain identification, a range of deep learning-based methods have been developed. For example, SpaGCN [20] leverages RGB channel data from areas surrounding the spots to gain histological insights, whereas stLearn [21], conST [22], and SiGra [23] employ various deep neural network models to extract image features. However, these approaches do not explicitly exploit the rich morphological details available in the images, such as cell locations and types, thereby offering limited insights into the interpretation of spatial domains. Recently, iIMPACT [24] has provided an approach to integrate the gene expression profile, spatial information, and detailed morphological information extracted from the artificial intelligence (AI)-reconstructed histology image from a single-sample SRT experiment. A summary of the existing clustering methods for analyzing SRT data is given in Table 1.

**Table 1.** Existing Methods applied to spatial domain identification in SRT data. Abbreviation: (i) Approach is either Bayesian (Bayes), Frequentist (Freq), or Deep Learning (DL); (ii) Dimension reduction technique is either Principal Component Analysis (PCA), Feature Selection (FS), AutoEncoder (AE), or Factor Analysis (FA). Methods included in the real data analysis are highlighted in bold.

| Method | Dimension Reduction Techniques | Tissue | Gene Expression | Spatial Profile | H&E Images | Approach | Language | Year |
| --- | --- | --- | --- | --- | --- | --- | --- | --- |
| **BASS** | PCA | Multi | √ | √ |  | Bayes | R/C++ | 2022 |
| BayesCafe | FS | Single | √ | √ |  | Bayes | R/C++ | 2023 |
| **BayesSpace** | PCA | Single | √ | √ |  | Bayes | R/C++ | 2021 |
| BNPSpace | FS | Single | √ | √ |  | Bayes | R/C++ | 2023 |
| CCST | AE | Single | √ | √ |  | DL | Python | 2022 |
| ConST | PCA | Single | √ | √ | √ | DL | Python | 2022 |
| **DR-SC** | FA | Single | √ | √ |  | Freq | R | 2022 |
| **iIMPACT** | PCA | Single | √ | √ | √ | Bayes | R/C++ | 2023 |
| **Louvain** | PCA | Single | √ |  |  | Freq | R | - |
| **Leiden** | PCA | Single | √ |  |  | Freq | Python | - |
| **SC-MEB** | PCA | Single | √ | √ |  | Freq | R/C++ | 2022 |
| SEDR | AE | Single | √ | √ |  | DL | Python | 2024 |
| **SpaGCN** | AE | Single | √ | √ | √ | DL | Python | 2021 |
| **STAGATE** | AE | Multi | √ | √ |  | DL | Python | 2022 |
| stLearn | PCA | Single | √ | √ | √ | Freq | Python | 2020 |

Nowadays, SRT studies often collect multiple adjacent sections from the same tissue or collect tissue samples from multiple individuals, while existing methods for analyzing a single tissue section could introduce bias to the clustering result. BASS [25] is the first statistical model tailored for spatial domain identification on multiple adjacent sections from the same tissue on single-cell SRT data. However, its performance is compromised when applied to multi-sample data from different individuals or when used on datasets generated by lower-resolution but widely used NGS-based technologies. Meanwhile, methods such as PASTE [26], STAGATE [27], DeepST [28], GraphST [29], and SPIRAL [30] only focus on the alignment of multi-sample SRT data by embedding the gene expression profiles to lower dimensional representations, yet the clustering step is not integrated and relies on the existing methods such as mclust [31] and Louvain [32]. A new class of methods, including IRIS [33] and SpaDo [34], offers enhanced clustering accuracy by performing spatial domain identification on multi-sample SRT data with single-cell reference dataset from the same tissue type. However, the effectiveness of these methods relies heavily on the availability of such a reference dataset at the single-cell level or annotation of cell types for each spatial location.

To address these challenges, we introduce BayeSMART (<u>Bayes</u>ian clustering of <u>m</u>ulti-sample sp<u>a</u>tially <u>r</u>esolved transcriptomics data), a fully Bayesian model designed for multi-sample SRT data to integrate molecular profiles, histological images, and spatial information to facilitate the spatial domain clustering when multiple sections or samples are available. Deep convolutional neural networks (CNN)-based methods such as H-DenseUNet [35], Micro-Net [36], Hover-Net [37], and HD-Yolo [38], have advanced histology image analysis by automating the segmentation, classification, and feature extraction of nuclei. We employed Hover-Net or HD-Yolo to generate the AI-reconstructed histology images from a multi-sample SRT experiment. This image profile at the single-cell level, together with the spot-level molecular profile and spatial information, was subsequently utilized by a Bayesian hierarchical finite mixture model to facilitate spatial domain identification. It is noteworthy that BayeSMART extends the capabilities of our previous work iIMPACT [24], which focuses on single-sample SRT datasets, by enabling multi-sample analysis and being applicable to a broader range of tissue types. We examined our method on two real datasets generated by NGS-based SRT technologies and found that it outperformed the existing approaches in terms of clustering accuracy and interpretability. To show that our method can also be applied to another type of SRT technologies with single-cell resolution, we analyzed a STARmap and a MERFISH dataset, and found that BayeSMART achieved better or similar performance on these datasets compared with the state-of-the-art methods, but with significantly less processing time.

## METHODS

In this section, we first define the multi-sample molecular, image, and geospatial profiles derived from NGS-based SRT data (e.g., ST and 10x Visium) and the paired AI-reconstructed histology images. For SRT data generated from those single-cell SRT technologies (e.g., STARmap and MERFISH), we employ a specialized approach as proposed by iIMPACT [24] to construct each of the three profiles (detailed in Supplementary Section S1). The results of the MERFISH data set are provided in Supplementary Section S2. The model achieves a better or similar performance on these datasets compared with the state-of-the-art methods but with a significantly shorter processing time (see details in Supplementary Section S3). We then outline the Bayesian statistical model that integrates the three profiles for multi-sample spatial clustering. The flowchart of the model is given in Figure 1. The detailed MCMC algorithms and posterior inference are presented in Supplementary Section S4. We evaluated the performance of the spatial domain identification using the widely adopted Adjusted Rand Index (ARI), which ranges from 0 to 1. Higher ARI values indicate greater similarity between the estimated spatial domain and the ground truth.

**Figure 1.**
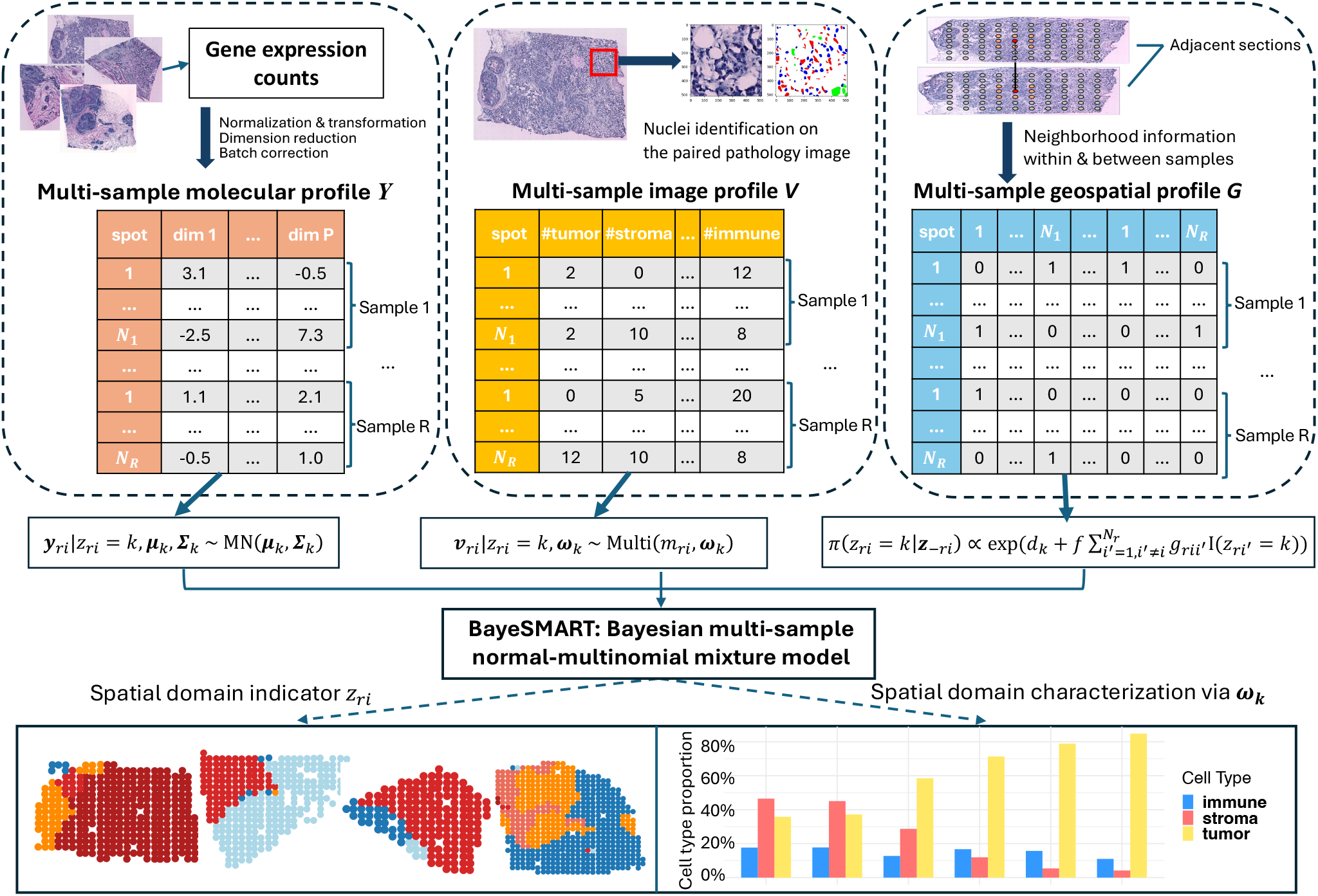
Flowchart of BayeSMART. BayeSMART performs multi-sample spatial domain identification analysis on SRT data. The gene expression count matrix from multiple samples are preprocessed to obtain the molecular profiles. Then AI-based methods are applied to each histology image of the samples to get the AI-reconstructed cell type information. Neighborhood information within and between samples is then utilized to construct the geospatial profile. With these three profiles prepared as matrices, a Bayesian normal-multinomial mixture model is employed to identify and interpret spatial domains across multiple samples.

### Data preparation

#### Multi-sample molecular profile ***Y***

Typically, a multi-sample SRT study measures the expression levels of overlapping gene sets from *R* samples (e.g., either adjacent sections from the same individual or samples from different individuals). The molecular profile of each sample *r* (*r* = 1, …, *R*) can then be denoted by an *N*_*r*_*× P*^common^ count matrix ***C***^(*r*)^, where *N*_*r*_ is the number of spots in sample *r, P*^common^ is the number of common genes shared by all the *R* samples, and each entry 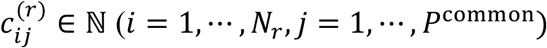 is the read count for gene *j* measured at spot *i* in sample *r*. To simplify the algebra, we combine the count matrices across all samples to an *N × P*^common^ count matrix 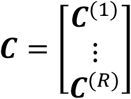, where 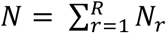 is the total number of spots in the study. Following many existing methods such as [39, 40], we apply library size normalization and a log transformation on ***C*** to conform to normality. As suggested by BASS [25], if the samples are adjacent sections from the same tissue, we apply SPARK-X [41] to first extract the top spatially variable genes (SVGs) out of *P*^common^ genes for each sample and then get the union of the SVGs among all samples to get *P*^select^ genes to proceed. If those multi-samples originate from distinct individuals, the above implementation may introduce additional noise due to the relatively small overlap among the identified SVG sets from different samples. In such a case, we select the top *P*^select^ = 2000 most highly variable genes (HVGs) in terms of their normalized gene expression levels as suggested by BayesSpace [19] and iIMPACT [24]. Detailed results and criteria of choosing between SVGs or HVGs are provided in Supplementary Section S5. After obtaining the *N* × *P*^select^matrix, we perform principal component analysis (PCA) for the dimension reduction, while alternative approaches such as t-SNE [42] or UMAP [43] are also applicable. We extract the top *P* principal components (PCs) and the resulting *N* × *P* matrix is denoted by 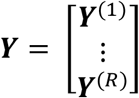, where each entry 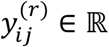 is the value of the *j*-th top PC at spot *i* in sample *r*. For simplicity of calculation, we follow BayesSpace [19] and iIMPACT [24] to choose *P* = 3, as the increase of *P* results in marginal or no improvements. Sensitivity analysis on the number of PCs is in Supplementary Figure S7. With the low dimensional representation matrix ***Y***, we finally perform the batch effect correction and alignment with Harmony [44] per recommendation [25, 45] and obtain the final form of the multi-sample molecular profile ***Y***.

#### Multi-sample image profile *V*

To integrate the multi-sample image profile into BayeSMART, we follow iIMPACT [24] and apply deep learning-based nuclei segmentation and classification algorithms to extract cellular abundance features from the paired histology images. To assess BayeSMART’s flexibility with various nuclei identification methods, we investigate our performance using two advanced methods, Hover-Net [37] and HD-Yolo [38]. Hover-Net is a CNN-based architecture pre-trained on various tissue types and can simultaneously segment and classify nuclei in multiple tissue types, including but not limited to breast cancer, colorectal adenocarcinoma, liver cancer, kidney cancer, and prostate cancer. HD-Yolo is another deep learning nuclei segmentation and classification approach specific to lung cancer, liver cancer, and breast cancer images. It is built upon the YOLO (You Only Look Once) architecture [46], designed for real-time object detection. Both methods provide the locations and types for all identified nuclei in the whole histology image. For the HER2-positive breast cancer dataset analyzed in the paper, we explored both methods, and the comparison is reported in the Supplementary Section S6. To match the molecular information measured at spots, which cover less than half the area (e.g., the area of all spots in 10x Visium platform is about 38% of the whole domain area), we count cells of different identified cell types within each spot and its expanded area (see Figure S20 in iIMPACT paper [24]). This ensures that all single-cell-level histology image information is utilized to enhance spatial clustering. For each sample *r*, the result can be summarized into an *N*_*r*_ × *Q* count matrix ***V***^(*r*)^, namely the cell type abundance table, where each entry 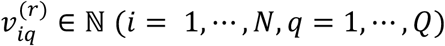 denotes the number of cells with type *q* observed at spot *i* and its expanded area in sample *r*.

There are situations where deep learning-based nuclei identification methods may not be appropriate, such as when a reliable pre-trained model is unavailable for a specific tissue type or the training histology images are missing or of poor quality. To ensure BayeSMART can be applied seamlessly, we can utilize an unsupervised, reference-free method called STdeconvolve [47]. This latent Dirichlet allocation (LDA)-based model estimates the relative cell type abundance of each spot in SRT data without a reference scRNA-seq dataset. LDA is a generative statistical model commonly used in natural language processing. STdeconvolve provides a built-in method to choose the optimal number of anonymous cell type *Q* given the SRT dataset. For sample *r*, the result of STdeconvolve can be summarized as an *N*_*r*_ × *Q* proportion matrix ***S***^(*r*)^, where each entry 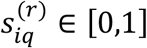 is the relative abundance of cell type *q* at spot *i* in sample *r*. To obtain the cell count matrix ***V***^(*r*)^, we multiply each entry of ***S***^(*r*)^ by a relatively large integer (e.g., 100) and round to the nearest integer to maintain the composition of cell types in each spot. We demonstrate that this special handling performed reasonably well in the real data analysis, although it sacrifices some interpretability of the resulting spatial domains.

### Multi-sample geospatial profile *G*

We represent the raw multi-sample SRT geospatial profile with an *N*_*r*_ × 2 matrix ***T***^(*r*)^, where each row 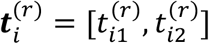 specifies the *x* and *y*-coordinates of the spot *i* in a two-dimensional Cartesian plane in sample *r*. Notably, spots—defined as the round areas for barcoded mRNA capture probes measuring gene expression—are arranged in square and triangular lattice grids for ST and 10x Visium platforms, respectively. Since the spots from the ST platform are squarely aligned, we consider the eight surrounding spots as neighbors for each non-boundary spot to capture more information from the neighborhood. This is achieved by identifying the eight closest spots to each non-boundary spot, typically including the spots directly above, below, left, right, and the four diagonal directions. For the 10x Visium platform, where spots are triangularly aligned, we consider the six surrounding spots as neighbors for each non-boundary spot. Moreover, when the samples are adjacent slides from the same tissue, neighborhood connections extend across two contiguous sections (detailed in the Supplementary Section S7). For all *N* spots across *R* samples, we can construct an *N* × *N* binary adjacent matrix ***G***, where an entry of one signifies neighboring spots, and zero indicates non-neighbors. The neighborhood structure ***G*** serves as our multi-sample geospatial profile, facilitating the integration of spatial data into the proposed Bayesian model.

### The proposed BayeSMART model

Our BayeSMART model, designed for spatial domain identification in a multi-sample SRT study, is an extension of iIMPACT [24], which focuses on analyzing single-sample SRT datasets. Given the multi-sample molecular and image profiles, we write the full data likelihood of our model as follows,

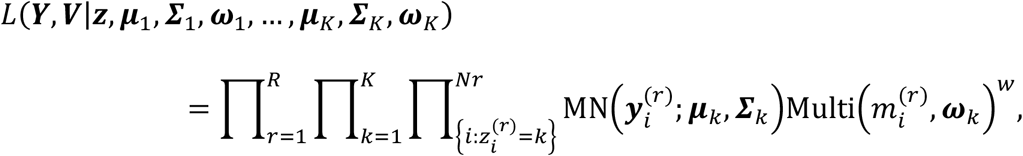

where 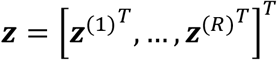 is the multi-sample spatial domain indicator vector, with 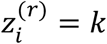 indicating that spot *i* in sample *r* belongs to spatial domain *k* shared by all samples. In the above equation, the first component assumes 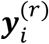 is from a multivariate normal (MN) distribution, where ***μ***_*k*_ and ***Σ***_*k*_ are the domain-specific mean vector and covariance matrix, respectively. The second component assumes 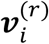 is from a multinomial distribution, where 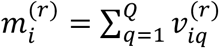 is the total number of cells observed within spot *i* in sample *r* (and if applicable, its expanded area) and ω_*k*_ represents the underlying relative abundance of cell types in spatial domain *k*, a key parameter that interprets and characterizes each identified spatial domain. The independence assumption between the two components is valid as the molecular profile ***Y*** and image profile ***V*** are generated from different sources. Note that the tuning parameter *w* ∈ [0,1] weighs the impact of the two profiles on the clustering process. Decreasing *w* parameterizes the multinomial likelihood towards one, thereby reducing the contribution of the image profile. We conduct a sensitivity analysis to determine for the best choice of *w*. The sensitivity analysis shows that choosing *w* within this range does not significantly affect the results (see Supplementary Section S8 for details). Specifically, our result suggests setting *w* = 0.1 for NGS-based SRT data, and for single-cell SRT data, a larger *w* ∈ [0.1, 0.5]. For the sake of computational efficiency (details in Supplementary Section S4), we adopt a conjugacy setting, which is ***μ***_*k*_|***Σ***_*k*_ ∼ MN(***ν***_0_, ***Σ***_*k*_/τ_0_), **Σ**_*k*_ ∼ IW(η_0_, **Φ**_0_), and ***ω***_*k*_ ∼ Dir(***α***_0_). Note that we generalize the prior setting for the normal component from the product normal-inverse-gamma distribution in iIMPACT [24] to the normal-inverse-Wishart (IW) distribution, eliminating constraints on the dimension reduction techniques used to generate the low-dimensional representation of the molecular profile.

In addition, we incorporate the geospatial information ***G*** by employing a Markov random field (MRF) prior [48, 49] on the multi-sample spatial domain indicator ***z*** as

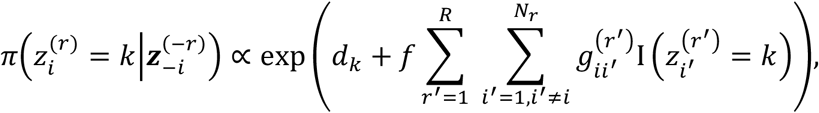

where 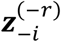 denotes the set of all entries in ***z*** among *R* samples excluding the *i*-th one in sample 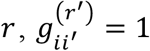 if spot *i* from sample *r* and spot *i′* from sample *r′* are neighbors, and 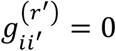 otherwise. The hyperparameters ***d*** = (*d*_1_, …, *d*_*K*_)^*T*^ ∈ ℝ^*K*^ control the number of spots belonging to each of the *K* spatial domains and *f* ∈ *ℝ*^+^controls the spatial smoothness. The purpose of using this prior is to encourage the neighboring spots to be clustered in the same spatial domain.

We recommend a weakly informative prior setting by choosing the MRF hyperparameter *d*_1_ = … = *d*_*k*_ = 1 and *f* = 1, the multivariate normal hyperparameters 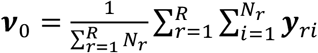 *τ*_0_ = 0.01, *η*_0_ = *P* + 1, and Φ_0_ = *I*_*P*×*P*_, and the multinomial hyperparameters *α*_01_ = … = *α*_0*Q*_ = 1. Given the assumption of independence 1) between the normal and multinomial components and 2) among the parameters of different spatial domains, and we use conjugate prior on all model parameters, *z, μ*_1_, …, *μ*_*k*_, Σ_1_, …, Σ_*k*_, *ω*_1_, …, *ω*_*k*_, their conditional distributions are all in closed form and easy to sample from. We can then apply Gibbs sampler, an MCMC algorithm for obtaining a sequence of observations approximated from a multivariate probability distribution. The details of the sampling process are in Supplementary Section S4.

### Choosing the number of spatial domains *K*

The number of spatial domains, *K*, can be determined using prior biological knowledge when available. In the absence of such information, the integrated completed likelihood (ICL) [50] can be used as a criterion for selecting *K*. The ICL is calculated by the following formula:

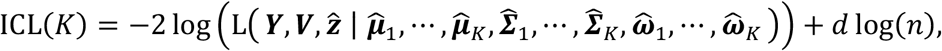

where 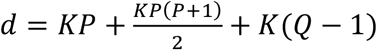 is the number of parameters in the model, and 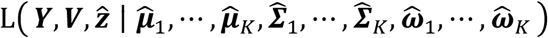 is complete data likelihood given by

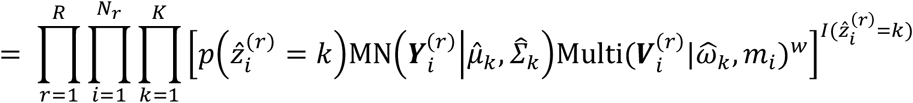

The results of the ICL analysis for each dataset and the ARIs corresponding to different values of *K* are presented in Figures S8 and S9 in the Supplementary Information, respectively. The results demonstrate that the ICL values corresponding to the number of manually annotated spatial domains are typically either the lowest or the second lowest. Additionally, the ARI values show considerable robustness across different choices of *K*.

## RESULTS

### Application to HER2-positive breast cancer ST dataset

Breast cancer is categorized into various subtypes, one of which is the HER2-positive classification. This subtype is characterized by a high level of HER2 (human epidermal growth factor receptor 2) expression in tumor cells [51, 52]. Approximately 15-2 0% of all breast cancer cases are HER2-positive. These tumors tend to grow aggressively and require intensive treatment [53, 54].

We applied BayeSMART to an ST dataset from an HER2-positive human breast cancer study [55]. Eight tumors were examined by three to six parallel sections under the original experiment, and only one section of each tumor was examined and annotated by a pathologist based on the morphology of the associated hematoxylin and eosin (H&E)-stained images (also known as pathology images, which are histology images from unhealthy tissues). Among the eight annotated sections, we selected sections A1, B1, C1, and H1 from four tumors to proceed with the following analysis (the quality control is provided in Supplementary Section S12). The H&E-stained images (Figure 2A) were annotated by a pathologist with six spatial domains: in situ cancer, invasive cancer, adipose tissue, breast glands, immune infiltrate, and connective tissue. Each sample contains 346, 295, 176, and 613 spots, respectively, and the number of common genes among these four samples is 13,2 91. After library-size normalization and log transformation, the top 2,000 HVGs were selected and used for the BayeSMART analysis.

**Figure 2.**
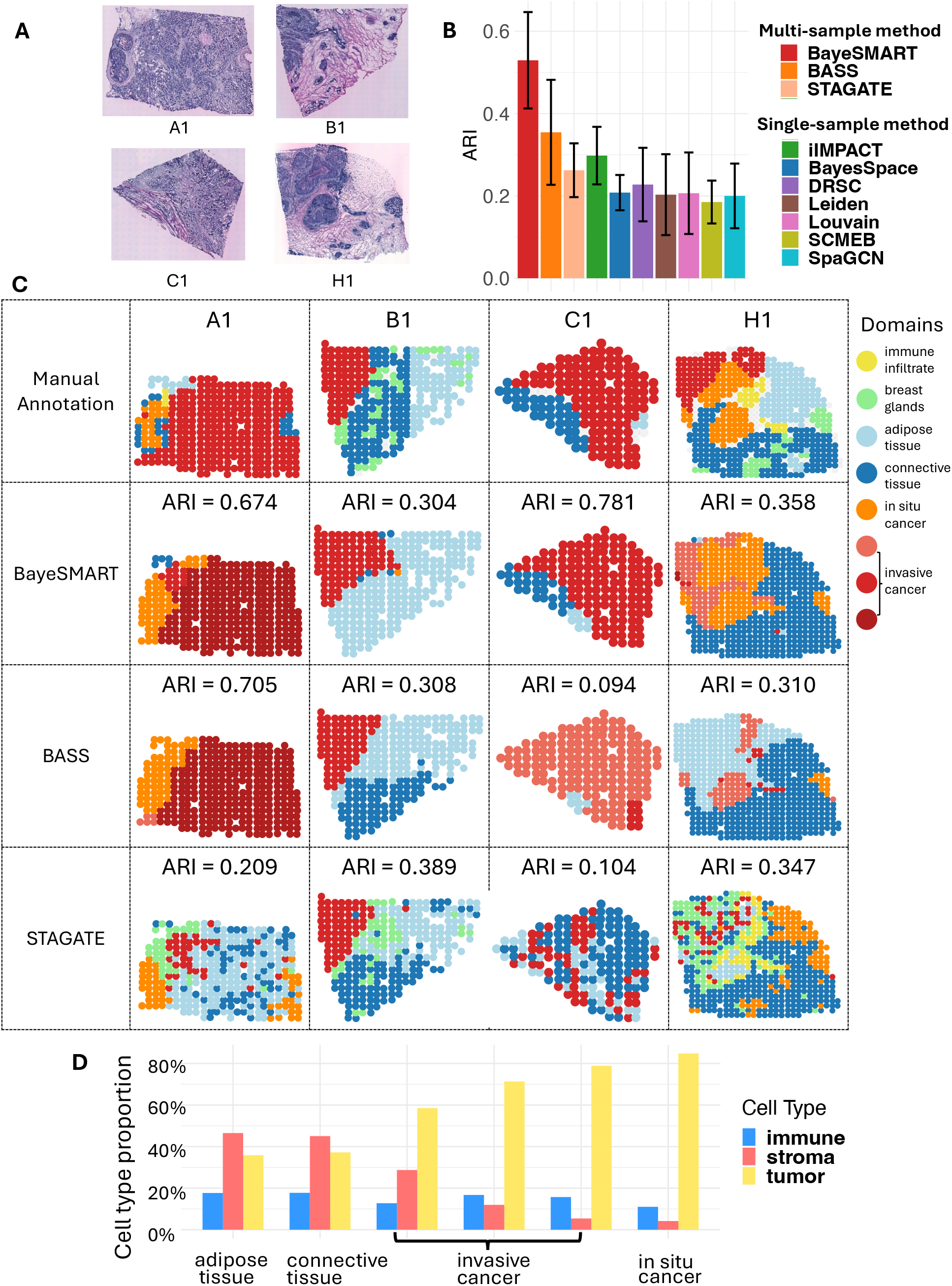
Result of the HER2-positive dataset. (**A**) H&E-stained images of four samples from breast cancer tumors of different individuals. (**B**) Barplots of ARI on four samples in ten compared approaches (BayeSMART, BASS, iIMPACT, BayeSpace, DRSC, Leiden, Louvain, SCMEB, SpaGCN, STAGATE). (**C**) The first row is the ground truth of the spatial domain annotated by a pathologist. The second, third, and fourth rows contain results for each section by BayeSMART, BASS, and STAGATE, respectively. For BayeSMART and BASS, the multi-sample analysis was applied. (**D**) Estimates (posterior means) of cell type proportion of the three cell types (observed in the AI-reconstructed histology image) on six domains.

First, we examined the performance of ten methods on this dataset. The implementation details of all the competing methods are given in Supplementary Section S11. As shown in Figure 2B, BayeSMART achieved the highest average ARI of 0.530 by detecting the spatial domains that closely resemble the ground truth. BASS is another method that can conduct multi-sample spatial domain identification and achieved the second-highest score of average ARI=0.354. STAGATE is a method based on an autoencoder for low-dimensional embedding that can be applied to simultaneously conduct clustering on multi-sample SRT datasets. Its performance is worse than that of Bayesian-based methods such as BayeSMART and BASS. All other methods (iIMPACT, BayesSpace, DRSC, Leiden, Louvain, SCMEB, and SpaGCN) can only deal with a single sample, thus for these methods we analyzed one sample at a time. As shown in Figure 2B, the performance of most of these methods is worse than that of multi-sample methods, BayeSMART, BASS, and STAGATE, with the performance of iIMPACT being better than that of STAGATE. Figure 2C provides the manual annotation (Row 1), and each sample’s spatial domain identification results for BayeSMART and BASS (Rows 2 and 3). Among these spatial domains, the cancer region (including invasive cancer and in situ cancer) is of great interest. When classifying the spatial domains as cancer and non-cancer regions, this identification task can also be treated as a binary classification problem, and we measured the performance with the area under the receiver operating characteristic curve (AUC), F1-score, and accuracy (ACC). BayeSMART reached the AUC, F1-score, and ACC of 0.832, 0.908, and 0.916, respectively, while BASS only reached 0.676, 0.815, and 0.852 (the details of the binary results are in Supplementary Section S9). It shows that BayeSMART performed the best when detecting cancer regions. It is worth noticing that for sample H1, even though the result of BASS reached an ARI of 0.310, it incorrectly classified the cancer region as a non-cancer region, which could introduce bias for the downstream analysis.

BayeSMART is characterized by its ability to interpret and define the detected spatial domains. Since the cell type information was AI-reconstructed, the Bayesian multinomial-normal mixture model can infer the relative abundance of each cell type in the spatial domains (Figure 2D). Other methods fall short of effectively integrating cell type information and directly interpreting these domains in a biologically meaningful manner. The posterior estimates of the cell type composition of each spatial domain are shown in Figure 2D. For example, the proportion of tumor cells in domains 3-6 is higher than the other domains, thus classified as cancer regions. This inference result is aligned with the cancer regions in the manual annotation, confirming that BayeSMART can capture correct biological information.

### Application to DLPFC 10x Visium dataset

We utilized BayeSMART to analyze the DLPFC data derived from the 10x Visium platform, which included 12 tissue sections from the dorsolateral prefrontal cortex of three adult donors. The DLPFC data encompass expression values for 33,538 genes across two pairs of tissue sections from three neurotypical adult donors. Each pair is composed of two 10 μm serial tissue sections, with the second pair located 300 μm posterior to the first, totaling 12 sections. Each section contains between 3,460 and 4,789 measured spots. Furthermore, we used manually annotated labels of seven laminar clusters, including six cortical layers (L1 to L6) and white matter (WM), as provided in the original publication, to serve as our ground truth for evaluating the performance of spatial domain detection [56]. The H&E-stained images of samples 151507-151510 from the first donor were exhibited in Figure 3A to assess the performance of BayeSMART.

**Figure 3.**
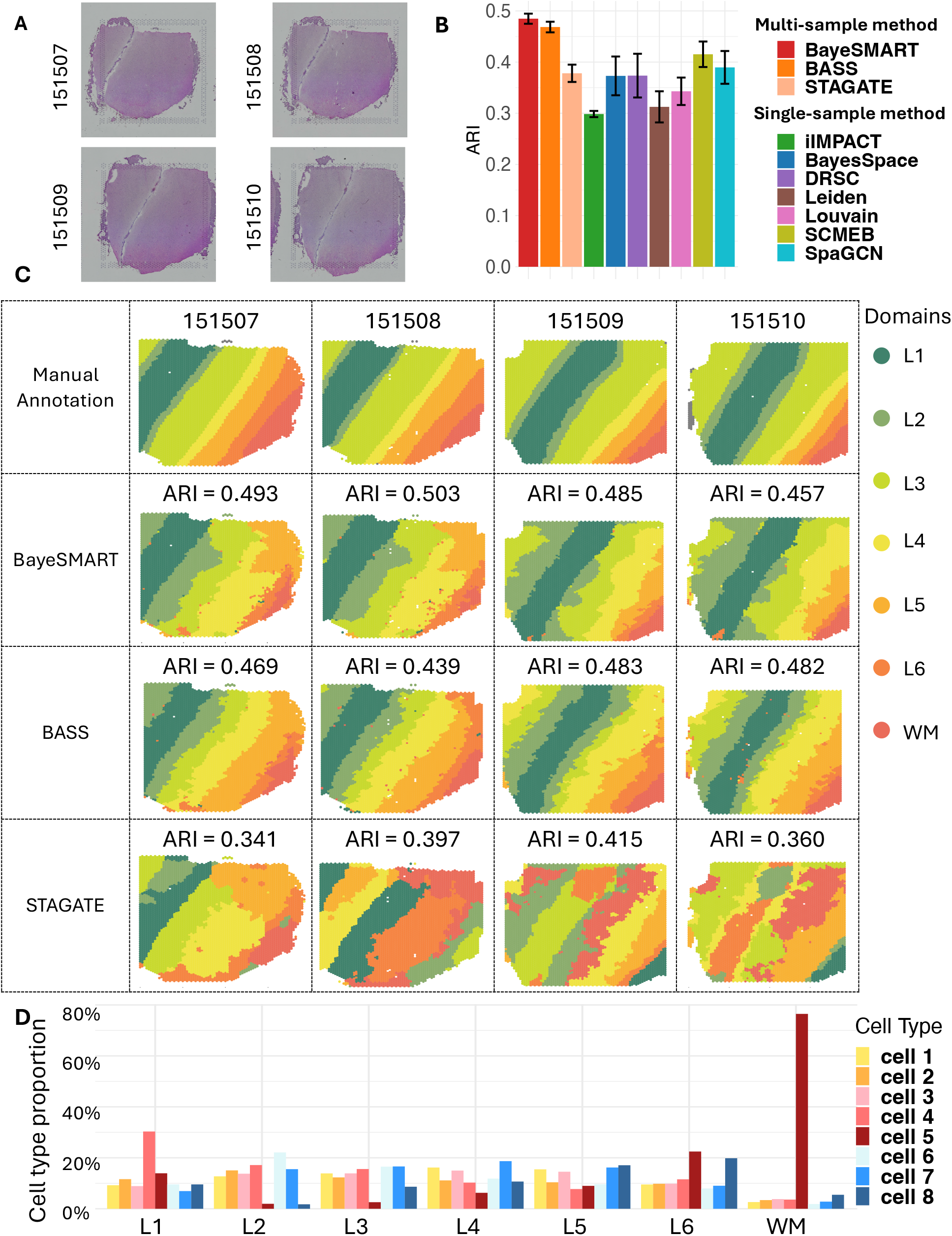
Result of the DLPFC dataset. (**A**) H&E-stained images of sections 151507, 151508, 151509, and 151510. (**B**) Barplots of ARI on four sections in ten compared approaches (BayeSMART, BASS, iIMPACT, BayeSpace, DRSC, Leiden, Louvain, SCMEB, SpaGCN, STAGATE). (**C**) The first row is the ground truth of the spatial domain annotated by pathologists. The second, third, and fourth rows contain results for each section by BayeSMART, BASS, and STAGATE, respectively. For BayeSMART and BASS, the multi-sample analysis was applied. (**D**) Estimates (posterior means) of cell type proportion of the eight “pseudo cell types” from STDeconvolve on seven domains.

Due to the absence of effective AI-based methods for nuclei segmentation and classification in the human dorsolateral prefrontal cortex, applying existing methods to this dataset would likely compromise the results of cell identification. Consequently, we have explored the cell information of the DLPFC using an unsupervised, reference-free method known as STdeconvolve (refer to the Method section for a detailed explanation of this technique). This method facilitates the determination of the optimal number of cell types, identified as eight for the DLPFC dataset.

STdeconvolve uses the original gene count matrix as input and predicts the cell composition for eight “pseudo cell types” at each spot. To align with BayeSMART’s requirement for an image profile matrix consisting of count data, we enhanced the resolution of the cell composition matrix by multiplying it by a relatively large integer (100 in our study) and rounding the results to the nearest integer to form our image profile. This approach offers a viable solution for situations where an image profile is unavailable or difficult to obtain when utilizing BayeSMART.

For the spatial information, since these four samples consist of adjacent sections from the same tissue, neighborhood information exists both within and between sections. We consider neighbors of each spot within the same section as well as those from adjacent section(s). Upon examining these four sections, a distinct parallel shift is observed in sections 151508 to 151510, using 151507 as the baseline. We shift sections 151508 to 151510 to the left by 4, 21, and 24 units, respectively. Then spots that share the same x- and y-axis values across adjacent sections are considered neighbors.

We compared BayeSMART with nine other methods, demonstrating that our model surpasses the others in performance (Figure 3B). Multi-sample methods BayeSMART and BASS show better performance in ARI than STAGATE and all the single-sample methods, highlighting the advantages of multi-sample Bayesian-based methods. STAGATE suffered from the label-switching problem across samples, challenging the interpretability of the clustering results. Specifically, BayeSMART achieved an average ARI of 0.485, outperforming the 0.468 achieved by BASS. The median improvement of BayeSMART over BASS across four samples is 5.12 %, and the p-value of the paired t-test between the resulting spatial domain labels by BayeSMART and BASS is less than 2.2 × 10^−16^, indicating that the results from BayeSMART are significantly different from those of BASS. We also applied BayeSMART on 12 slides from three donors together, and the result shows the robustness of BayeSMART on multiple samples from different individuals (Supplementary Section S10).

For the interpretation of spatial domain identification results, since STDeconvolve was applied, the cell types in the image profile matrix are “pseudo cell types” denoted by cell types 1 to 8. STDeconvolve also provides the most common genes for each “cell type,” which provides a way of interpreting each “pseudo cell type.” As shown in Figure 3D, domain L1 is dominated by cell type 4, and domain WM is characterized by cell type 5.

### Application to STARmap dataset

To demonstrate that BayeSMART is also applicable to data from single-cell resolution SRT platforms, we applied BayeSMART to a STARmap dataset. This dataset originates from the mouse medial prefrontal cortex (mPFC), comprising three tissue sections from the mPFC of different mice. The mPFC is a critical region in the frontal lobe’s anterior portion, pivotal in executing high-level cognitive functions such as decision-making, memory, attention, and emotion [57]. Structurally, the mPFC consists of four layers: L1, L2/3, L5, and L6. It predominantly contains excitatory pyramidal neurons (approximately 80-90%) and inhibitory GABAergic interneurons (about 10-2 0%), which play key roles in orchestrating cortical network dynamics and maintaining connections with distant targets [25, 58]. The tissue sections, labeled as BZ5 (1,049 cells), BZ9 (1,053 cells), and BZ14 (1,088 cells), were analyzed for gene expression across a consistent set of 166 genes. The annotation of four distinct cortical layers—L1, L2/3, L5, and L6—for each single cell was curated from [25]. Since STARmap achieves single-cell resolution, AI-based methods are not needed in this scenario. To facilitate model execution, we constructed “pseudo spots” by overlaying squarely ordered grids on the sample (see Figure 4A). These three samples are adjacent sections from the same tissue, so neighborhood information exists between two contiguous sections. For the geospatial profile, we considered the neighbors of each spot both within the same section and across sections to enhance performance. We first compared BayeSMART with the other nine methods on this dataset (Figure 4B). BayeSMART achieved the highest average ARI of 0.775 across three samples, while BASS reached an average ARI of 0.771. The paired t-test comparing the resulting spatial domain labels of each single cell from BayeSMART and BASS yielded a p-value of 2.1 × 10^−6^, indicating that BayeSMART’s performance is significantly better than that of BASS. Additionally, iIMPACT also delivered commendable results by analyzing each sample separately, achieving an average ARI of 0.723. The other seven methods—BayesSpace, DRSC, Leiden, Louvain, SCMEB, SpaGCN, and STAGATE—yielded significantly lower ARI scores. Figure 4C displays the detailed spatial domain identification results for each sample by BayeSMART, BASS, and STAGATE. The plot illustrates that the layers L5 and L6 identified by BayeSMART are cleaner and smoother, whereas the corresponding layers identified by BASS exhibit some noise. For sample BZ9, BayeSMART achieved an improvement of 13.91% over BASS.

**Figure 4.**
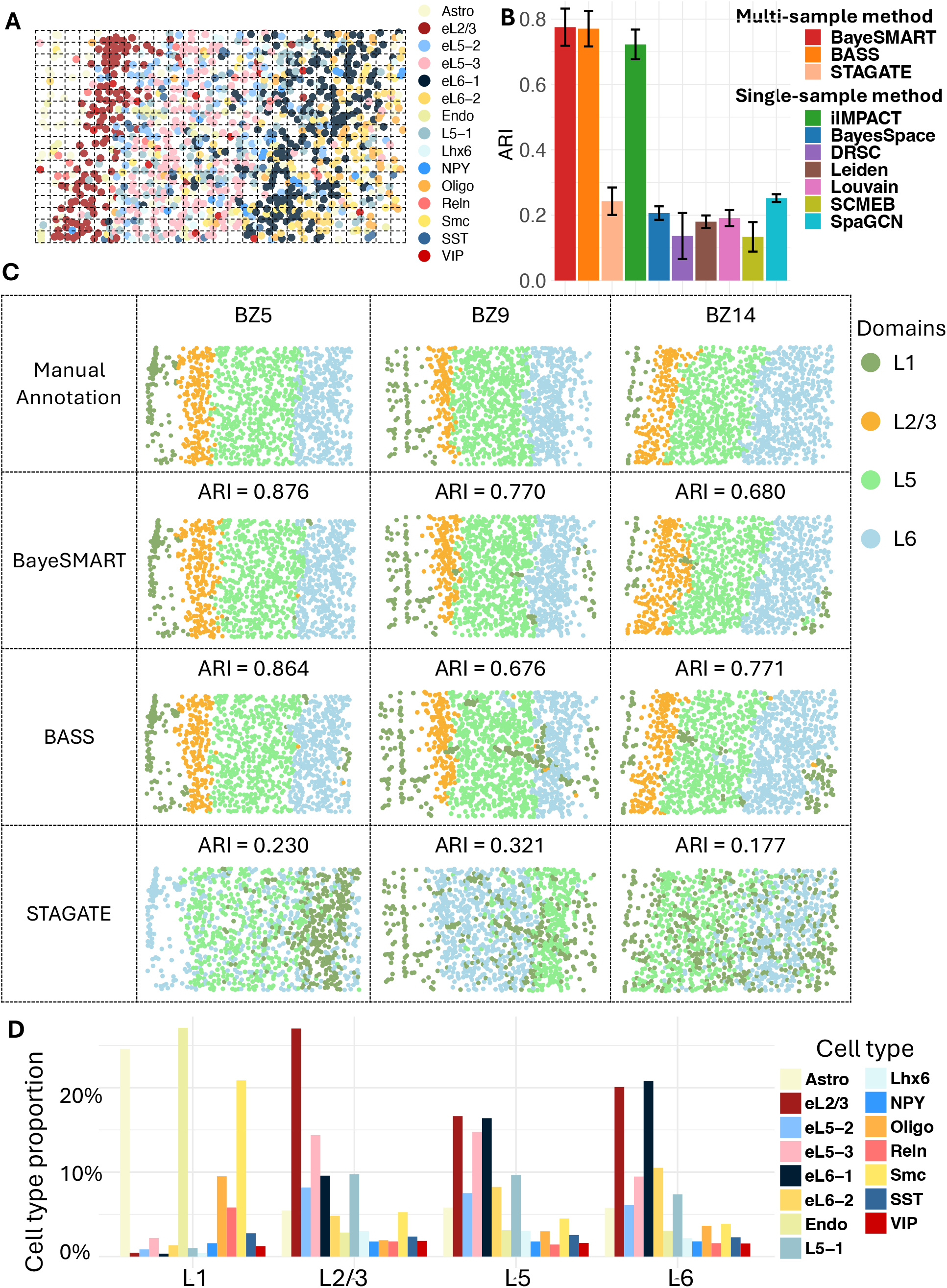
Result of the STARmap dataset. (**A**) Section BZ14 with cells annotated by cell types. An example of a grid (spot) assignment is given. (**B**) Barplots of ARI on four sections in ten compared approaches (BayeSMART, BASS, iIMPACT, BayeSpace, DRSC, Leiden, Louvain, SCMEB, SpaGCN, STAGATE). (**C**) The first row is the annotated layer structure of the tissue sections from the original study. The second, third, and fourth rows contain results for each section by BayeSMART, BASS, and STAGATE, respectively. For BayeSMART and BASS, the multi-sample analysis was applied. (**D**) Estimates (posterior means) of cell type proportion of the 15 cell types on four domains.

For the STARmap dataset, the analysis and understanding of the spatial domain identification results are based on the cell-type labels provided by the original study. The posterior estimates of the cell type composition for each spatial domain are presented in Figure 4D. Specifically, Layer L1 is characterized by higher levels of Astrocytes and Endothelial cells, and Layer L2/3 contains the highest concentration of early L2/3 neurons. Layer L5 predominantly features early L2/3, eL5-3, and eL6-1 neurons, while Layer L6 is marked by a significant presence of early L2/3 and eL6-1 neurons.

## DISCUSSION

This paper presents BayeSMART, a Bayesian model that advances multi-sample SRT analysis by integrating molecular, image, and spatial profiles from NGS-based SRT experiments. By analyzing adjacent sections or samples from different individuals, BayeSMART mitigates biases inherent in single-section studies, offering a more comprehensive analysis. Unlike BASS, BayeSMART also leverages neighborhood information between adjacent sections from the same tissue, enhancing its robustness. In addition, the application of BayeSMART to diverse technologies like ST, 10x Visium, STARmap, and MERFISH has demonstrated its versatility and superior performance in identifying spatial domains across different platforms and tissue types. Moreover, the utilization of AI for histology image reconstruction, coupled with spatial context, has demonstrated substantial improvements in the identification of spatial domains, as supported by our extensive case studies. Our exploration of deep learning-based image feature extraction methods, such as Hover-Net and HD-Yolo, and our application of the reference-free method STDeconvolve in scenarios where images are unavailable or unsuitable for analysis further enhance the broad applicability of our approach. Moreover, BayeSMART’s interpretability of the detected spatial domains provides deeper biological insights from spatial clustering results, and therefore could improve our understanding of spatial heterogeneity in healthy and diseased tissues. Despite these advances, certain limitations must be acknowledged. The performance of BayeSMART is influenced by the quality of the histology images, and therefore relies on the accuracy of pre-trained deep learning algorithms for image processing. Next, since the reference-free method STDeconvolve is not designed for multi-sample analysis, we recommend using STDeconvolve only on adjacent slides when a suitable deep learning method is not available to generate the image profile, rather than on samples from different individuals. Furthermore, while BayeSMART can provide insights into the cellular composition of tissues, the biological interpretation of spatial domains remains a challenge and warrants further investigation.

In conclusion, BayeSMART stands as a significant contribution to the field of SRT, provides a robust framework for integrating complex datasets, and reveals meaningful biological insights. Future studies should focus on applying BayeSMART to a broader range of tissue types and pathological conditions. Also, refining nuclei segmentation and classification methods for histology images may further enhance BayeSMART’s performance. Next, the best setting of the tuning parameter *w*, which adjusts the weight of the image profiles, can be further determined for different SRT platforms and not limited to the ones we examined in this study. Moreover, different batch-effect correction and alignment methods can be explored on various SRT datasets to further enhance the performance. Lastly, the MRF prior in BayeSMART currently only incorporates binary neighborhood information. We may further generalize it to utilize both local and global spatial information for each spot.

### Key Points

- BayeSMART represents the first multi-sample spatial domain identification method tailored for NGS-based SRT data.
- The method enhances spatial domain identification’s accuracy and computational efficiency by integrating cell-type abundance information derived from AI-reconstructed histological images.
- It provides interpretability for clustering results through the estimation of posterior cell-type compositions within each identified spatial domain.
- The major existing methods for spatial domain identification were compared with BayeSMART using datasets from various tissue types and experimental platforms.

## AUTHOR CONTRIBUTIONS STATEMENT

YG carried out the research, analyzed the data, interpreted the result, and was the major contributor to writing the manuscript. CB summarized and ran competing methods (BayesSpace, DRSC, Leiden, Louvain, SCMEB, SpaGCN, and STAGATE). CT conducted the image analysis with Hover-Net. RR conducted the image analysis with HD-Yolo. YM provided guidance on the experiment with reference-free version of IRIS. GX, LX and QL conceived the study and supervised the statistical modeling and analyses. All authors read and approved the final manuscript.

## DATA AVAILABILITY

The code and data of BayeSMART are accessible through the GitHub repository at https://github.com/yg2485/BayeSMART. The implementation of Hover-Net can be found at https://github.com/vqdang/hover_net?tab=readme-ov-file, and the implementation of HD-Yolo can be found at https://github.com/impromptuRong/hd_wsi. The tutorial of STDeconvolve is provided at https://jef.works/STdeconvolve/.

## Reference

1. Burgess DJ. Spatial transcriptomics coming of age, Nature Reviews Genetics 2019;20:317–317.

2. Zhang M, Sheffield T, Zhan X et al. Spatial molecular profiling: platforms, applications and analysis tools, Briefings in Bioinformatics 2021;22.

3. Moor AE, Itzkovitz S. Spatial transcriptomics: paving the way for tissue-level systems biology, Curr Opin Biotechnol 2017;46:126–133.

4. Ståhl PL, Salmén F, Vickovic S et al. Visualization and analysis of gene expression in tissue sections by spatial transcriptomics, Science 2016;353:78–82.

5. Rodriques SG, Stickels RR, Goeva A et al. Slide-seq: A scalable technology for measuring genome-wide expression at high spatial resolution, Science 2019;363:1463–1467.

6. Stickels RR, Murray E, Kumar P et al. Highly sensitive spatial transcriptomics at near-cellular resolution with Slide-seqV2, Nat Biotechnol 2021;39:313–319.

7. Vickovic S, Eraslan G, Salmén F et al. High-definition spatial transcriptomics for in situ tissue profiling, Nat Methods 2019;16:987–990.

8. Lubeck E, Coskun AF, Zhiyentayev T et al. Single-cell in situ RNA profiling by sequential hybridization. Nat Methods. 2014, 360–361.

9. Chen KH, Boettiger AN, Moffitt JR et al. RNA imaging. Spatially resolved, highly multiplexed RNA profiling in single cells, Science 2015;348:aaa6090.

10. Wang X, Allen WE, Wright MA et al. Three-dimensional intact-tissue sequencing of single-cell transcriptional states, Science 2018;361.

11. Larsson L, Frisén J, Lundeberg J. Spatially resolved transcriptomics adds a new dimension to genomics, Nature Methods 2021;18:15–18.

12. Asp M, Bergenstrahle J, Lundeberg J. Spatially Resolved Transcriptomes-Next Generation Tools for Tissue Exploration, Bioessays 2020;42:e1900221.

13. Lee J, Yoo M, Choi J. Recent advances in spatially resolved transcriptomics: challenges and opportunities, BMB Reports 2022;55:113–124.

14. Moses L, Pachter L. Museum of spatial transcriptomics, Nat Methods 2022;19:534–546.

15. Thrane K, Eriksson H, Maaskola J et al. Spatially Resolved Transcriptomics Enables Dissection of Genetic Heterogeneity in Stage III Cutaneous Malignant Melanoma, Cancer Res 2018;78:5970–5979.

16. Satija R, Farrell JA, Gennert D et al. Spatial reconstruction of single-cell gene expression data, Nat Biotechnol 2015;33:495–502.

17. Kiselev VY, Andrews TS, Hemberg M. Challenges in unsupervised clustering of single-cell RNA-seq data, Nat Rev Genet 2019;20:273–282.

18. Zhu Q, Shah S, Dries R et al. Identification of spatially associated subpopulations by combining scRNAseq and sequential fluorescence in situ hybridization data, Nat Biotechnol 2018.

19. Zhao E, Stone MR, Ren X et al. Spatial transcriptomics at subspot resolution with BayesSpace, Nat Biotechnol 2021;39:1375–1384.

20. Hu J, Li X, Coleman K et al. SpaGCN: Integrating gene expression, spatial location and histology to identify spatial domains and spatially variable genes by graph convolutional network, Nature Methods 2021;18:1342–1351.

21. Pham D, Tan X, Xu J et al. stLearn: integrating spatial location, tissue morphology and gene expression to find cell types, cell-cell interactions and spatial trajectories within undissociated tissues. Cold Spring Harbor Laboratory, 2020.

22. Zong Y, Yu T, Wang X et al. conST: an interpretable multi-modal contrastive learning framework for spatial transcriptomics. Cold Spring Harbor Laboratory, 2022.

23. Tang Z, Li Z, Hou T et al. SiGra: single-cell spatial elucidation through an image-augmented graph transformer, Nature Communications 2023;14.

24. Jiang X, Wang S, Guo L et al. Integrating Image and Molecular Profiles for Spatial Transcriptomics Analysis. Cold Spring Harbor Laboratory, 2023.

25. Li Z, Zhou X. BASS: multi-scale and multi-sample analysis enables accurate cell type clustering and spatial domain detection in spatial transcriptomic studies, Genome Biology 2022;23.

26. Zeira R, Land M, Strzalkowski A et al. Alignment and integration of spatial transcriptomics data, Nature Methods 2022;19:567–575.

27. Dong K, Zhang S. Deciphering spatial domains from spatially resolved transcriptomics with an adaptive graph attention auto-encoder, Nature Communications 2022;13.

28. Xu C, Jin X, Wei S et al. DeepST: identifying spatial domains in spatial transcriptomics by deep learning, Nucleic Acids Research 2022;50:e131–e131.

29. Long Y, Ang KS, Li M et al. Spatially informed clustering, integration, and deconvolution of spatial transcriptomics with GraphST, Nature Communications 2023;14.

30. Guo T, Yuan Z, Pan Y et al. SPIRAL: integrating and aligning spatially resolved transcriptomics data across different experiments, conditions, and technologies, Genome Biology 2023;24.

31. Fraley C, Raftery AE. mclust: Gaussian Mixture Modelling for Model-Based Clustering, Classification, and Density Estimation. Comprehensive R Archive Network (CRAN).

32. Blondel VD, Guillaume J-L, Lambiotte R et al. Fast unfolding of communities in large networks, Journal of Statistical Mechanics: Theory and Experiment 2008;2008:P10008.

33. Ma Y, Zhou X. Accurate and efficient integrative reference-informed spatial domain detection for spatial transcriptomics, Nature Methods 2024:1–14.

34. Duan B, Chen S, Cheng X et al. Multi-slice spatial transcriptome domain analysis with SpaDo, Genome Biology 2024;25:73.

35. Li X, Chen H, Qi X et al. H-DenseUNet: hybrid densely connected UNet for liver and tumor segmentation from CT volumes, IEEE transactions on medical imaging 2018;37:2663–2674.

36. Raza SEA, Cheung L, Shaban M et al. Micro-Net: A unified model for segmentation of various objects in microscopy images, Med Image Anal 2019;52:160–173.

37. Graham S, Vu QD, Raza SEA et al. Hover-Net: Simultaneous segmentation and classification of nuclei in multi-tissue histology images, Med Image Anal 2019;58:101563.

38. Rong R, Sheng H, Jin KW et al. A Deep Learning Approach for Histology-Based Nucleus Segmentation and Tumor Microenvironment Characterization, Modern Pathology 2023;36:100196.

39. L Lun AT, Bach K, Marioni JC. Pooling across cells to normalize single-cell RNA sequencing data with many zero counts, Genome Biology 2016;17.

40. McCarthy DJ, Campbell KR, Lun ATL et al. Scater: pre-processing, quality control, normalization and visualization of single-cell RNA-seq data in R, Bioinformatics 2017;33:1179–1186.

41. Zhu J, Sun S, Zhou X. SPARK-X: non-parametric modeling enables scalable and robust detection of spatial expression patterns for large spatial transcriptomic studies, Genome Biology 2021;22.

42. Van der Maaten L, Hinton G. Visualizing data using t-SNE, Journal of machine learning research 2008;9.

43. Becht E, McInnes L, Healy J et al. Dimensionality reduction for visualizing single-cell data using UMAP, Nature biotechnology 2019;37:38–44.

44. Korsunsky I, Millard N, Fan J et al. Fast, sensitive and accurate integration of single-cell data with Harmony, Nature Methods 2019;16:1289–1296.

45. Tran HTN, Ang KS, Chevrier M et al. A benchmark of batch-effect correction methods for single-cell RNA sequencing data, Genome Biology 2020;21.

46. Redmon J, Divvala S, Girshick R et al. You Only Look Once: Unified, Real-Time Object Detection. IEEE.

47. Miller BF, Huang F, Atta L et al. Reference-free cell type deconvolution of multi-cellular pixel-resolution spatially resolved transcriptomics data, Nature Communications 2022;13.

48. Clifford P. Markov Random Fields in Statistics. Disorder in Physical Systems: A Volume in ∼Honour of John M. Hammersley. Oxford University Press, 1990, 19–32.

49. Morris R, Descombes X, Zerubia J. Fully Bayesian image segmentation-an engineering perspective. IEEE Comput. Soc.

50. Biernacki C, Celeux G, Govaert G. Assessing a mixture model for clustering with the integrated completed likelihood, IEEE transactions on pattern analysis and machine intelligence 2000;22:719–725.

51. Wang J, Xu B. Targeted therapeutic options and future perspectives for HER2-positive breast cancer, Signal Transduction and Targeted Therapy 2019;4.

52. Mano MS, Rosa DD, Azambuja ED et al. The 17q12-q21 amplicon: Her2 and topoisomerase-IIalpha and their importance to the biology of solid tumours, Cancer Treat Rev 2007;33:64–77.

53. Fitzmaurice C, Abate D, Abbasi N et al. Global, Regional, and National Cancer Incidence, Mortality, Years of Life Lost, Years Lived With Disability, and Disability-Adjusted Life-Years for 29 Cancer Groups, 1990 to 2017, JAMA Oncology 2019;5:1749.

54. Sharma P. Major Strides in HER2 Blockade for Metastatic Breast Cancer, N Engl J Med 2020;382:669–671.

55. Andersson A, Larsson L, Stenbeck L et al. Spatial deconvolution of HER2-positive breast cancer delineates tumor-associated cell type interactions, Nature Communications 2021;12.

56. Maynard KR, Collado-Torres L, Weber LM et al. Transcriptome-scale spatial gene expression in the human dorsolateral prefrontal cortex, Nature Neuroscience 2021;24:425–436.

57. Carlén M. What constitutes the prefrontal cortex?, Science 2017;358:478–482.

58. Xu P, Chen A, Li Y et al. Medial prefrontal cortex in neurological diseases, Physiological Genomics 2019;51:432–442.

