## Supplementary Information for "BayeSMART: Bayesian Clustering of Multi-sample Spatially Resolved Transcriptomics Data"

### Supplementary Information
### for
### BayeSMART: Bayesian Clustering of Multi-sample Spatially Resolved Transcriptomics Data

#### S1. Special Handling of Multi-sample SRT Data at the Single-cell Resolution

Imaging-based and some new sequencing-based SRT techniques offer higher resolution than traditional NGS-based techniques, which capture the molecular profile at a single-cell level. Data from these platforms typically provide the spatial distribution and types of cells on the tissue section from the original study. To adapt BayeSMART to single-cell SRT data from techniques such as STARmap and MERFISH, we manually add a square lattice grid of appropriate size to the entire domain and consider each square unit as a spot. Note that these "pseudo spots" fill the entire domain. Thus, there is no gap between two adjacent spots, and each cell can be assigned to exactly one spot. The non-boundary spots each have eight neighboring spots within the sample, and potentially two neighbors from the adjacent sections above and below. The geospatial profile $\boldsymbol{G}$ is constructed by considering both the neighboring spots within the same section and between adjacent sections. The definition of neighbors for each spot within the slide is shown in Figure S6, and the neighborhood information between two adjacent slides is further discussed in Section S7. To construct the molecular profile $\boldsymbol{Y}$, we first average the counts of each gene across all cells within each spot. Library-size normalization, log transformation, and dimension reduction steps are then applied to the dataset in the same way as for traditional NGS-based techniques. For the "image" profile $\boldsymbol{V}$, we directly count the cells of different types in each spot.

#### S2. Application to Mouse Hypothalamus MERFISH Dataset

MERFISH is another single-cell resolution SRT technology. In this paper, adjacent tissue sections from the preoptic region of the mouse hypothalamus were analyzed. This region is crucial within the brain's center, containing multiple nuclei that regulate a wide range of social behaviors, such as reproduction and circadian rhythms, as well as homeostatic functions like neuroendocrine and cardiovascular regulation. Three sections—Bregma-0.14 (5,926 cells), Bregma-0.19 (5,803 cells), and Bregma-0.24 (5,543 cells)—are under investigation in this study, with gene expression measurements collected for a common set of 155 genes. The cells in each tissue section were carefully annotated into eight spatial domains: the third ventricle (V3), bed nuclei of the stria terminalis (BST), columns of the fornix (fx), medial preoptic area (MPA), medial preoptic nucleus (MPN), periventricular hypothalamic nucleus (PV), paraventricular hypothalamic nucleus (PVH), and paraventricular nucleus of the thalamus (PVT).

Since MERFISH also achieves single-cell resolution, AI-based methods are not required for this dataset. To facilitate model execution, we constructed "pseudo spots" by overlaying squarely ordered grids on the sample (see Figure S1A). Given that these three samples are adjacent sections from the same tissue, neighborhood information also exists between two contiguous sections. For the multi-sample geospatial profile, we considered the neighbors of each spot both within the same section and across sections to enhance performance. We first compared BayeSMART with the other nine methods on this dataset (Figure S1B). BayeSMART and BASS achieved significantly better performance than other single-sample methods. BASS slightly outperformed BayeSMART with an average ARI of 0.556 among three samples, where BayeSMART achieved 0.500. STAGATE also reached an average ARI of 0.423. The other seven methods—BayesSpace, DRSC, iIMPACT, Leiden, Louvain, SCMEB, and SpaGCN—yielded ARI scores below 0.3. These results show the effectiveness of multi-sample SRT spatial domain identification methods compared with single sample methods. Figure S1C displays the detailed spatial domain identification results for each sample by BayeSMART and BASS. Since the V3 region is a narrow line structure, and to be able to run BayeSMART we need to assign grids to the sample, which is coarser than V3, the corresponding domain detected by BayeSMART is wider than the ground truth.

The posterior estimates of the cell type composition for each spatial domain are presented in Figure S1D. Specifically, the domain BST, PV, and MPN are characterized by higher levels of inhibitory cells, and PVT contains the highest concentration of excitatory cells. MPA has higher levels of both inhibitory and pericytes.

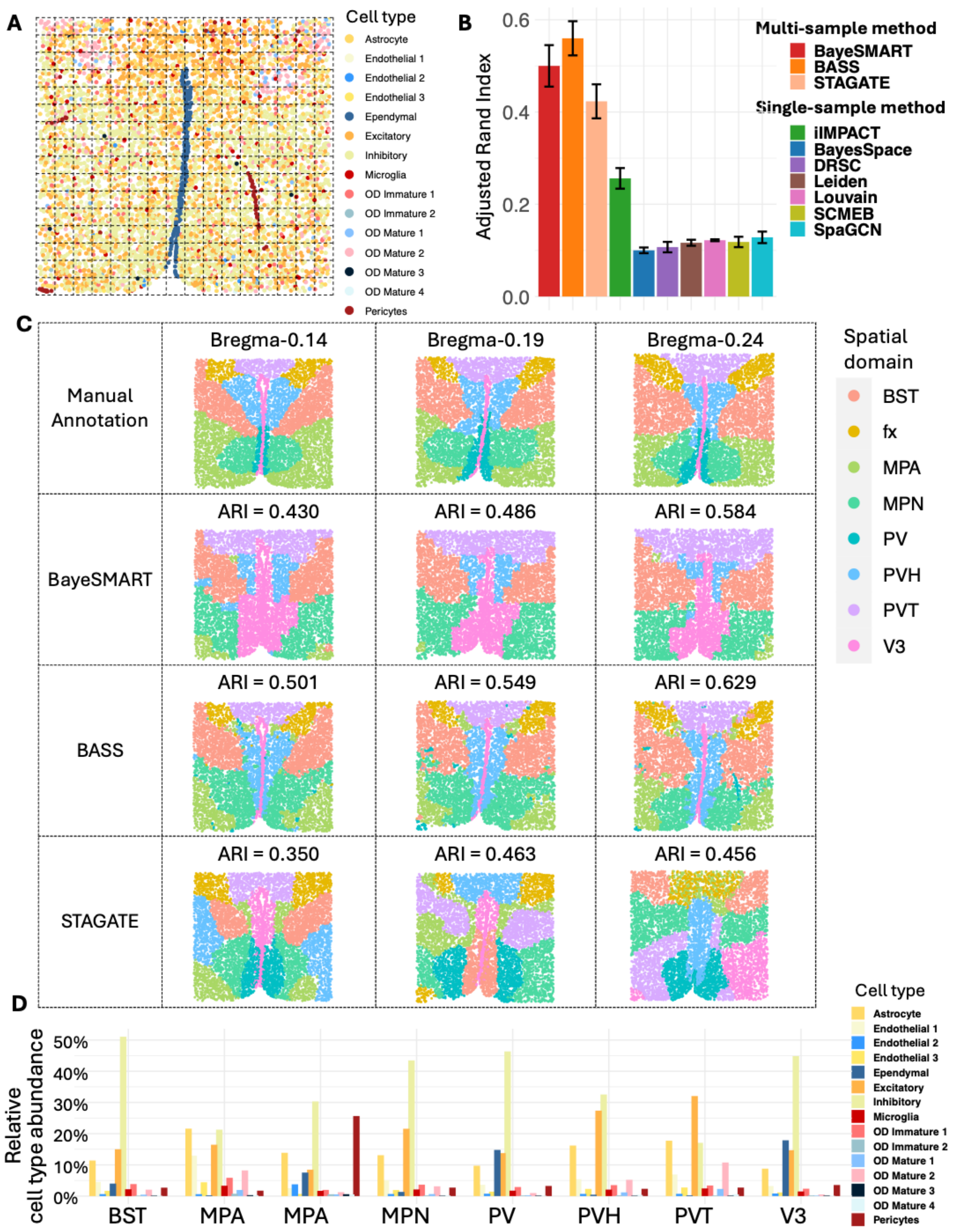


Figure S1. Results of the MERFISH dataset. (**A**) Section Bregma-0.24 with cells annotated by cell types. An example of grid (pseudo spot) assignment is given. (**B**) Barplots of ARI for three sections in ten compared approaches (BayeSMART, BASS, iIMPACT, BayeSpace, DRSC, Leiden, Louvain, SCMEB, SpaGCN, STAGATE). (**C**) The first row shows the manual annotation of tissue sections from the original study. The second, third, and fourth rows contain results for each section by BayeSMART, BASS, and STAGATE, respectively. (**D**) Estimates (posterior means) of relative cell type abundance for the 15 cell types on eight domains.

**S3. Running Time of BayeSMART, BASS, and iIMPACT on the MERFISH Dataset**

The MERFISH dataset includes 12 samples available for processing, thus making it suitable for evaluating the scalability of various methods. We compared the processing times of BayeSMART, iIMPACT, and BASS on 3, 6, and 12 samples to assess the scalability of the methods. BayeSMART, a multi-sample version improved from iIMPACT, can process the multi-sample spatial domain identification in a shorter amount of time. In contrast, BASS requires more than ten times the processing duration of BayeSMART.

| | 3 samples | 6 samples | 12 samples |
|---|---|---|---|
| BayeSMART | 1.027 mins | 1.905 mins | 3.685 mins |
| iIMPACT | 1.778 mins | 3.214 mins | 7.315mins |
| BASS | 11.495 mins | 22.737 mins | 44.709 mins |

Table S1. The processing time of BayeSMART, iIMPACT, and BASS on 3, 6, and 12 samples.

#### S4. Details of the MCMC Algorithm

The full data likelihood of the proposed Bayesian finite normal-multinomial mixture model is given as follows,

$$\begin{aligned}
&L(\boldsymbol{Y}, \boldsymbol{V} | \boldsymbol{z}, \boldsymbol{\mu}_1, \dots, \boldsymbol{\mu}_K, \boldsymbol{\Sigma}_1, \dots, \boldsymbol{\Sigma}_K, \boldsymbol{\omega}_1, \dots, \boldsymbol{\omega}_K) \\
=& \prod_{r=1}^{R} \prod_{k=1}^{K} \prod_{i=1}^{N_r} \mathrm{I}(z_i^{(r)} = k) f(\boldsymbol{y}_i^{(r)}, \boldsymbol{v}_i^{(r)} | z_i^{(r)} = k, \boldsymbol{\mu}_k, \boldsymbol{\Sigma}_k, \boldsymbol{\omega}_k),
\end{aligned}$$

where $R$ is the total number of samples, $K$ is the pre-defined number of clusters, $N_r$ is the number of spots in the $r$th sample, $\mathrm{I}(\cdot)$ denotes the indicator function, and

$$\begin{aligned}
&L(\boldsymbol{y}_i^{(r)}, \boldsymbol{v}_i^{(r)} | z_i^{(r)} = k, \boldsymbol{\mu}_k, \boldsymbol{\Sigma}_k, \boldsymbol{\omega}_k) \\
&= L(\boldsymbol{y}_i^{(r)} | z_i^{(r)} = k, \boldsymbol{\mu}_k, \boldsymbol{\Sigma}_k) f(\boldsymbol{v}_i^{(r)} | z_i^{(r)} = k, \boldsymbol{\omega}_k)^w \\
&= \mathrm{MN}(\boldsymbol{y}_i^{(r)}; \boldsymbol{\mu}_k, \boldsymbol{\Sigma}_k) \mathrm{Multi}(\boldsymbol{v}_i^{(r)}; m_i^{(r)}, \boldsymbol{\omega}_k)^w \\
&= (2\pi)^{-P/2} |\boldsymbol{\Sigma}_k|^{-1/2} \exp\left(-\frac{1}{2}(\boldsymbol{y}_i^{(r)} - \boldsymbol{\mu}_k)^\top \boldsymbol{\Sigma}_k^{-1} (\boldsymbol{y}_i^{(r)} - \boldsymbol{\mu}_k)\right) \left(\frac{m_i!}{\prod_{q=1}^{Q} v_{iq}^{(r)}!} \prod_{q=1}^{Q} \omega_{kq}^{v_{iq}^{(r)}}\right)^w \\
&\propto |\boldsymbol{\Sigma}_k|^{-1/2} \exp\left(-\frac{1}{2}(\boldsymbol{y}_i^{(r)} - \boldsymbol{\mu}_k)^\top \boldsymbol{\Sigma}_k^{-1} (\boldsymbol{y}_i^{(r)} - \boldsymbol{\mu}_k)\right) \left(\prod_{q=1}^{Q} \omega_{kq}^{v_{iq}^{(r)}}\right)^w.
\end{aligned}$$

We assume an independent prior structure 1) between the normal and multinomial subcomponents, and 2) among the parameters belonging to different spatial domains. Thus, the joint distribution of priors for parameters can be written as

$$\begin{aligned}
&\pi(\boldsymbol{z}, \boldsymbol{\mu}_1, \dots, \boldsymbol{\mu}_K, \boldsymbol{\Sigma}_1, \dots, \boldsymbol{\Sigma}_K, \boldsymbol{\omega}_1, \dots, \boldsymbol{\omega}_K) \\
&= \pi(\boldsymbol{z}) \pi(\boldsymbol{\mu}_1, \dots, \boldsymbol{\mu}_K, \boldsymbol{\Sigma}_1, \dots, \boldsymbol{\Sigma}_K) \pi(\boldsymbol{\omega}_1, \dots, \boldsymbol{\omega}_K)^w \\
&= \pi(\boldsymbol{z}) \prod_{k=1}^{K} \pi(\boldsymbol{\mu}_k, \boldsymbol{\Sigma}_k) \prod_{k=1}^{K} \pi(\boldsymbol{\omega}_k)^w \\
&= \pi(\boldsymbol{z}) \prod_{k=1}^{K} \pi(\boldsymbol{\mu}_k | \boldsymbol{\Sigma}_k) \pi(\boldsymbol{\Sigma}_k) \prod_{k=1}^{K} \pi(\boldsymbol{\omega}_k)^w.
\end{aligned}$$

We assign the Markov random field (MRF) prior to the spatial domain indicator $\boldsymbol{z}$ as

$$\pi(z_i^{(r)} = k|\boldsymbol{z}_{-i}^{(-r)}) \propto \exp\left(d_k + f\sum_{i'=1,i'\neq i}^{N_r} g_{ii'}^{(r)}\mathrm{I}(z_{i'}^{(r)} = k)\right),$$

We assign the conjugate priors to other parameters, listed as follows, so that the Gibbs sampler can be applied for posterior sampling.

$$\begin{aligned}
&\boldsymbol{\mu}_k|\boldsymbol{\Sigma}_k \sim \mathrm{MN}(\boldsymbol{\nu}_0, \boldsymbol{\Sigma}/\tau_0)\\
&\qquad\text{or equivalently, } \pi(\boldsymbol{\mu}_k|\boldsymbol{\Sigma}_k) = (2\pi/\tau_0)^{-P/2}|\boldsymbol{\Phi}_0|^{-1/2}\exp\left(-\frac{\tau_0}{2}(\boldsymbol{\mu}_k - \boldsymbol{\nu}_0)^\top\boldsymbol{\Phi}_0^{-1}(\boldsymbol{\mu}_k - \boldsymbol{\nu}_0)\right),\\
&\quad\boldsymbol{\Sigma}_k \sim \mathrm{IW}(\eta_0, \boldsymbol{\Phi}_0)\\
&\qquad\text{or equivalently, } \pi(\boldsymbol{\Sigma}_k) = \frac{|\boldsymbol{\Phi}_0|^{\eta_0/2}}{2^{\eta_0 P/2}\Gamma_P(\eta_0/2)}|\boldsymbol{\Sigma}_k|^{-(\eta_0+P+1)/2}\exp\left(-\frac{1}{2}\mathrm{tr}\left(\boldsymbol{\Phi}_0\boldsymbol{\Sigma}_k^{-1}\right)\right),
\end{aligned}$$

and

$$\boldsymbol{\omega}_k \sim \mathrm{Dir}(\boldsymbol{\alpha}_0) \text{ or equivalently, } \pi(\boldsymbol{\omega}_k) = \frac{\Gamma\left(\sum_{q=1}^{Q}\alpha_{0q}\right)}{\prod_{q=1}^{Q}\Gamma(\alpha_{0q})}\prod_{q=1}^{Q}\omega_{kq}^{\alpha_{0q}-1},$$

where $\Gamma_P(\cdot)$ and $\Gamma(\cdot)$ denote the $P$-dimensional and univariate gamma function.

We recommend a weakly informative prior setting by choosing the MRF hyperparameters $d_1 = \ldots = d_K = 1$ and $f = 1$, the multivariate normal hyperparameters $\boldsymbol{\nu}_0 = \frac{1}{\sum_{r=1}^{R} N_r}\sum_{r=1}^{R}\sum_{i=1}^{N_r}\boldsymbol{y}_{ri}$, $\tau_0 = 0.01$, $\eta_0 = P + 1$, and $\boldsymbol{\Phi}_0 = \boldsymbol{I}_{P\times P}$ (i.e., the $P$-by-$P$ identity matrix), and the multinomial hyperparameters $\alpha_{01} = \ldots = \alpha_{0Q} = 1$.

The full posterior distribution of the proposed Bayesian normal-multinomial mixture model is given in the following formula.

$$\begin{aligned}
&\pi(\boldsymbol{z}, \boldsymbol{\mu}_1, \ldots, \boldsymbol{\mu}_K, \boldsymbol{\Sigma}_1, \ldots, \boldsymbol{\Sigma}_K, \boldsymbol{\omega}_1, \ldots, \boldsymbol{\omega}_K|\boldsymbol{Y}, \boldsymbol{V}) \propto\\
&L(\boldsymbol{Y}, \boldsymbol{V}|\boldsymbol{z}, \boldsymbol{\mu}_1, \ldots, \boldsymbol{\mu}_K, \boldsymbol{\Sigma}_1, \ldots, \boldsymbol{\Sigma}_K, \boldsymbol{\omega}_1, \ldots, \boldsymbol{\omega}_K)\pi(\boldsymbol{z}, \boldsymbol{\mu}_1, \ldots, \boldsymbol{\mu}_K, \boldsymbol{\Sigma}_1, \ldots, \boldsymbol{\Sigma}_K, \boldsymbol{\omega}_1, \ldots, \boldsymbol{\omega}_K)
\end{aligned}$$

Posterior sampling is employed by the MCMC algorithm. Our primary interest lies in identifying spatial domains and the interactive zone *via* inferring the spatial domain indicator vector $\boldsymbol{z}$, and in characterizing domain-specific relative abundance of cell types *via* inference of $\boldsymbol{\omega}_1, \ldots, \boldsymbol{\omega}_K$. Since we use conjugate priors on all model parameters, $\boldsymbol{z}, \boldsymbol{\mu}_1, \ldots, \boldsymbol{\mu}_K, \Sigma_1, \ldots, \Sigma_K$, $\boldsymbol{\omega}_1, \ldots, \boldsymbol{\omega}_K$, their conditional distributions are all in closed form and easy to sample from. Consequently, we can rely on the Gibbs sampler, an MCMC algorithm for obtaining a sequence of observations approximated from a multivariate probability distribution when direct sampling is difficult. To be specific, we perform the following steps sequentially at each MCMC iteration after a random initialization.

**Update the spatial domain indicator $\boldsymbol{z}$**: We update $z_1^{(r)}, \ldots, z_{N_r}^{(r)}$ sequentially for each of the $R$ samples. To allocate spot $i$ in the $r$th sample to one of the $K$ spatial domains, we sample $z_i$ from a multinomial distribution,

$$z_i^{(r)}|\cdot \sim \text{Multi}(1, (\pi(z_i^{(r)} = 1|\cdot)/e, \ldots, \pi(z_i^{(r)} = K|\cdot)/e)),$$

where

$$\begin{aligned}
\pi(z_i^{(r)} = k|\cdot) &\propto L(\boldsymbol{y}_i^{(r)}, \boldsymbol{v}_i^{(r)}|z_i^{(r)} = k, \boldsymbol{\mu}_k, \boldsymbol{\Sigma}_k, \boldsymbol{\omega}_k)\pi(z_i^{(r)} = k|\boldsymbol{z}_{-i}^{(-r)}) \\
&\propto |\boldsymbol{\Sigma}_k|^{-1/2} \exp\left(-\frac{1}{2}(\boldsymbol{y}_i^{(r)} - \boldsymbol{\mu}_k)^\top \boldsymbol{\Sigma}_k^{-1}(\boldsymbol{y}_i^{(r)} - \boldsymbol{\mu}_k)\right) \left(\prod_{q=1}^{Q} \omega_{k,q}^{v_{iq}^{(r)}}\right)^w \\
&\quad \exp\left(d_k + f \sum_{i'=1, i' \neq i}^{N_r} g_{ii'}^{(r)} \mathrm{I}(z_{i'}^{(r)} = k)\right)
\end{aligned}$$

and the normalization constant $e = \sum_{k=1}^{K} \pi(z_i^{(r)} = k|\cdot)$.

**Update the domain-specific relative abundance of cell types $\boldsymbol{\omega}_k$'s**: We update $\boldsymbol{\omega}_1, \ldots, \boldsymbol{\omega}_K$ sequentially. For each spatial domain $k$, we draw a sample of $\boldsymbol{\omega}_k$ from a Dirichlet distribution,

$$\boldsymbol{\omega}_k|\cdot \sim \text{Dir}(\boldsymbol{\alpha}_k),$$

where the concentration parameters $\boldsymbol{\alpha}_k = (\alpha_{k1}, \ldots, \alpha_{kQ})$ with each entry $\alpha_{kq} = \alpha_{0q} + \sum_{r=1}^{R} \sum_{i=1}^{N_r} \mathrm{I}(z_i^{(r)} =$

$k)v_{iq}^{(r)}$. Note that the last term $\sum_{r=1}^{R}\sum_{i=1}^{N_r}\mathrm{I}(z_i^{(r)}=k)v_{iq}^{(r)}$ denotes the total number of cells with type $q$ observed in spatial domain $k$.

**Update the domain-specific low-dimensional representation of gene expression mean $\boldsymbol{\mu}_k$'s:** We update $\boldsymbol{\mu}_1,\ldots,\boldsymbol{\mu}_K$ sequentially. For each spatial domain $k$, we draw a sample of $\boldsymbol{\mu}_k$ from a multivariate normal distribution,

$$\boldsymbol{\mu}_k|\cdot \sim \mathrm{MN}(\boldsymbol{\nu}_k, \boldsymbol{\Sigma}_k/\tau_k),$$

where $\boldsymbol{\nu}_k=(\boldsymbol{\nu}_0\tau_0+n_k\bar{\boldsymbol{y}}_k)/(\tau_0+n_k)$ and $\tau_k=\tau_0+n_k$. Note that $n_k=\sum_{r=1}^{R}\sum_{i=1}^{N_r}\mathrm{I}(z_i^{(r)}=k)$ is the number of spots allocated to spatial domain $k$ and $\bar{\boldsymbol{y}}_k=\frac{1}{n_k}\sum_{r=1}^{R}\sum_{i=1}^{N_r}\mathrm{I}(z_i^{(r)}=k)\boldsymbol{y}_i^{(r)}$ denotes the average low-dimensional gene expression value over all the spots allocated to spatial domain $k$. If PCA is chosen to reduce the dimension of the SRT molecular profile, then we can further set $\Sigma_k$ to a $P$-by-$P$ diagonal matrix due to orthogonality among principal components. In this special case, we can draw each entry in $\boldsymbol{\mu}_k$ independently,

$$\mu_{kj}|\cdot \sim \mathrm{N}(\nu_{kj}, \sigma_{kj}^2/\tau_k),$$

where $\nu_{kj}=(\nu_{0j}\tau_0+n_k\bar{y}_{kj})/(\tau_0+n_k)$.

**Update the domain-specific covariance matrix of the low-dimensional representation of gene expression $\boldsymbol{\Sigma}_k$'s:** We update $\boldsymbol{\Sigma}_1,\ldots,\boldsymbol{\Sigma}_K$ sequentially. For each spatial domain $k$, we draw a sample of $\boldsymbol{\Sigma}_k$ from an inverse-Wishart distribution,

$$\boldsymbol{\Sigma}_k|\cdot \sim \mathrm{IW}(\eta_k, \boldsymbol{\Phi}_k),$$

where $\eta_k=\eta_0+n_k$ and $\boldsymbol{\Phi}_k=\boldsymbol{\Phi}_0+\sum_{r=1}^{R}\sum_{i=1}^{N_r}\mathrm{I}(z_i^{(r)}=k)(\boldsymbol{y}_i^{(r)}-\bar{\boldsymbol{y}}_k)(\boldsymbol{y}_i^{(r)}-\bar{\boldsymbol{y}}_k)^\top+\frac{\tau_0 n_k}{\tau_0+n_k}(\bar{\boldsymbol{y}}_k-\boldsymbol{\nu}_0)(\bar{\boldsymbol{y}}_k-\boldsymbol{\nu}_0)^\top$.

#### S5. The Choice between SVGs and HVGs for Generating Multi-sample Molecular Profile

To demonstrate that the choice between using SVGs or HVGs can enhance clustering results across different datasets, we evaluated their performance on the HER2-positive breast cancer dataset and the DLPFC dataset. Three gene selection methods were applied to each dataset, and their efficacy was compared. As illustrated in Table S2, gene selection notably improved the performance of BayeSMART on both datasets. Specifically, for the HER2-positive breast cancer dataset, SVGs and HVGs achieved average ARI scores of 0.473 and 0.530, respectively. However, a paired t-test between these results yielded a p-value of 0.617, suggesting no significant difference in the performance of the two gene selection methods. The Venn diagram in Figure S2 shows that selecting 2000 SVGs per sample resulted in 309 overlapping genes, representing 15.45% overlap. In contrast, for the DLPFC dataset, 880 genes overlapped out of 2000 (44% overlap), and SVGs resulted in a slightly higher average ARI of 0.485 compared to 0.474 with HVGs, with a p-value of 0.05 from the paired t-test, indicating an insignificance in difference. To optimize performance, we established a criterion for selecting gene selection methods based on a 30% overlap threshold. When the overlap of SVGs selected from each sample exceeds 30%, SVGs are preferable; otherwise, HVGs should be used.

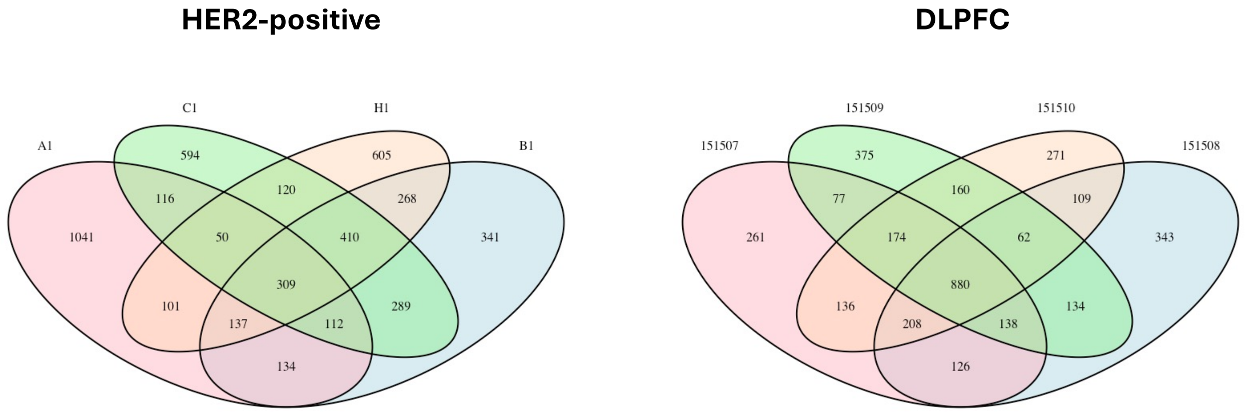


Figure S2. Venn diagram of SVGs selected by SPARK-X for each sample of HER2-positive dataset (left) and DLPFC dataset (right).

| HER2-positive | | | | | |
|---|---|---|---|---|---|
| | A1 | B1 | C1 | H1 | Average |
| All genes | 0.686 | 0.260 | 0.227 | 0.376 | 0.386 |
| SVGs | 0.697 | 0.379 | 0.425 | 0.392 | 0.473 |
| HVGs | 0.674 | 0.304 | 0.781 | 0.358 | **0.530** |
| DLPFC | | | | | |
| | 151507 | 151508 | 151509 | 151510 | Average |
| All genes | 0.447 | 0.468 | 0.418 | 0.389 | 0.431 |
| SVGs | 0.493 | 0.503 | 0.485 | 0.457 | **0.485** |
| HVGs | 0.483 | 0.483 | 0.476 | 0.453 | 0.474 |

Table S2. ARI of BayeSMART on two datasets using different gene selection methods. When all genes are used, no subset of genes is pre-selected for subsequent data processing. We applied SPARK-X for selecting the SVGs, and the scran package for selecting the HVGs.

**S6. Comparison of Hover-Net and HD-Yolo on the HER2-positive Breast Cancer Dataset**

Hover-Net and HD-Yolo are two deep learning methods applicable to breast cancer datasets, facilitating the segmentation and classification of nuclei within tissue images. They yield information on cell types and the spatial coordinates of each identified cell. Conversely, STDeconvolve, an unsupervised approach, takes ST gene counts as input to yield the cellular composition at each spot. We employed four approaches to analyze the HER2-positive breast cancer dataset to generate the image profile: 1) Hover-Net for both cell segmentation and classification, 2) HD-Yolo for both cell segmentation and classification, 3) Hover-Net for cell segmentation coupled with STDeconvolve for cell classification, and 4) HD-Yolo for cell segmentation coupled with STDeconvolve for cell classification. In the last two approaches, after localizing each cell and assigning them to specific spots, the number of cells in each spot is obtained and STDeconvolve is then used to get the composition and count of each cell type per spot.

| | A1 | B1 | C1 | H1 | Average |
|---|---|---|---|---|---|
| Hover-Net | 0.685 | 0.305 | 0.798 | 0.332 | **0.530** |
| HD-Yolo | 0.658 | 0.347 | 0.394 | 0.015 | 0.353 |
| Hover-Net + STDeconvolve | 0.664 | 0 | 0 | 0.272 | 0.234 |
| HD-Yolo + STDeconvolve | 0.664 | 0.273 | 0.305 | 0.268 | 0.378 |

Table S3. Results from BayeSMART on the HER2-positive breast cancer dataset using four approaches to generate the image profile.

#### S7. Multi-sample Geospatial Profile $G$ for Adjacent Sections

When working with multiple samples from different individuals, neighborhood information is considered only within each sample. However, when the samples are adjacent slides from the same tissue, neighborhood connections are extended across the two contiguous sections. Take the DLPFC dataset as an example: since samples 151507-151510 consist of adjacent sections from the same tissue, we consider neighbors of each spot within the same section as well as those from adjacent section(s). Without extra information, the general strategy is to treat the spots at the same position on the probe from two adjacent slides as neighbors. For example, on the 10x Visium platform, the spot at position '10×12' on slides 151507 and 151508 both correspond to the barcoded area at the 10th row and 12th column.

However, for slides 151507-151510, we observed from the H&E-stained images a clear parallel shift in the positioning of the tissue on the Visium slides. For illustration purposes, we provided the shift between 151507 and 151508 as an example in Figure S3A. 151509 and 151510 can be adjusted in a similar manner. Given that the distance between slides 151507 and 151508, as well as between 151509 and 151510, is 10 μm, and the distance between slides 151508 and 151509 is 300 μm (only three times the distance between spots), which are much smaller than the size of the whole slide, we considered a parallel shift by moving slides 151508-151510 closer to 151507 by 4, 21, and 24 spots along the x-axis, respectively. Then, spots that share the same x- and y-axis values across adjacent sections are considered neighbors. Figure S3B presents the BayeSMART results for slides 151507-151510 using these two strategies. Incorporating this additional spatial information into the geospatial profile resulted in a minor improvement. However, in the absence of specific information about the relative positions of adjacent slides, treating spots with the same position on two adjacent slides as neighbors generally provides a reliable result.

For the STARmap and MERFISH datasets, since we assign the same squarely aligned grids to each section, the grids with same x- and y-grid assignments from two adjacent sections are considered as neighbors. We integrated this neighborhood information into the geospatial profile $\boldsymbol{G}$ along with neighborhood information within each section.

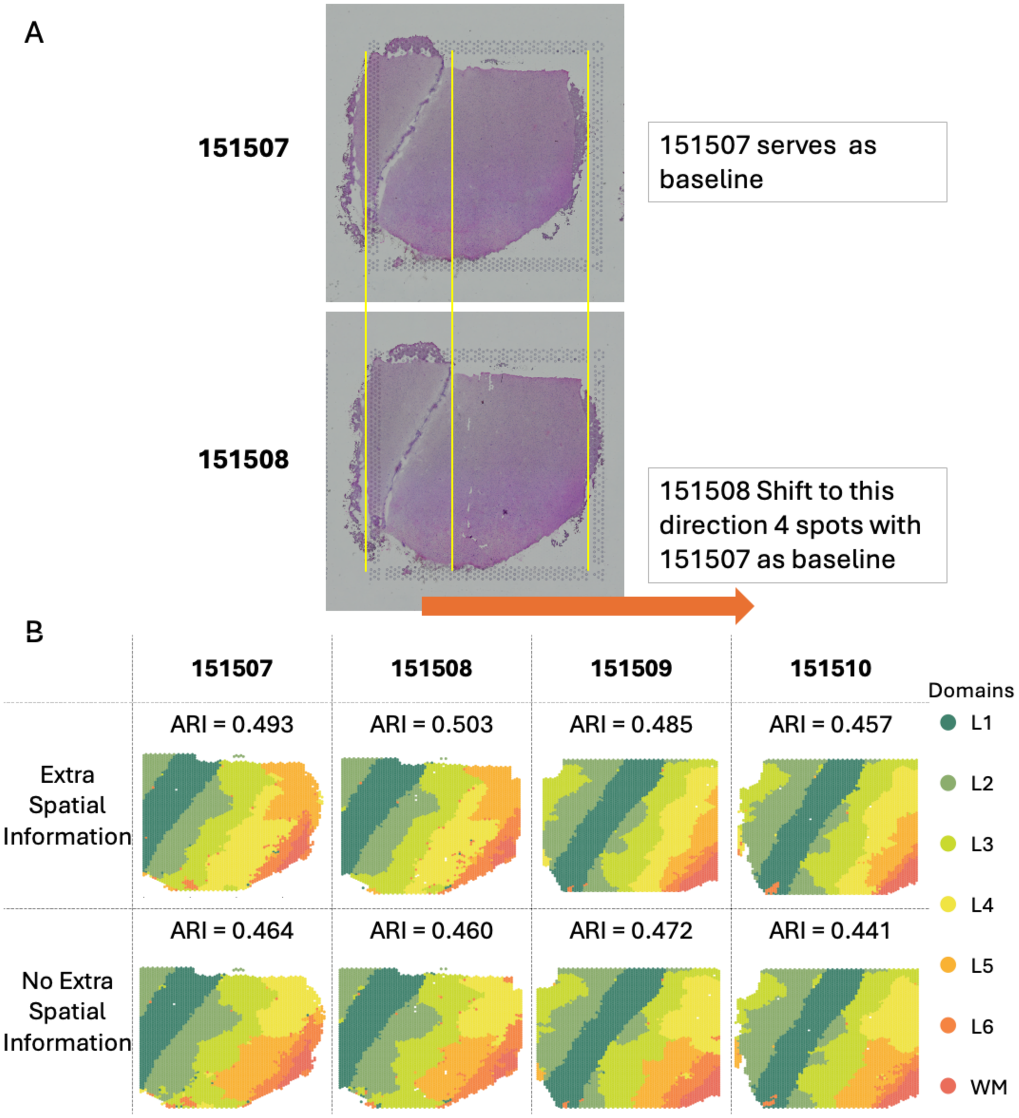


Figure S3. Neighborhood information between adjacent slides of DLPFC. (A) The tissue slide 151508 was positioned 4 spots lower relative to slide 151507, which serves as the baseline. (B) BayeSMART results for slides 151507-151510. The first row shows the outcomes when additional spatial information is incorporated into the geospatial profile $\boldsymbol{G}$, while the second row presents the results when such spatial information is unavailable. In this case, we consider two spots with identical positions on adjacent slides as neighbors.

#### S8. The Choice of Multi-sample Image Profile Weight *w*

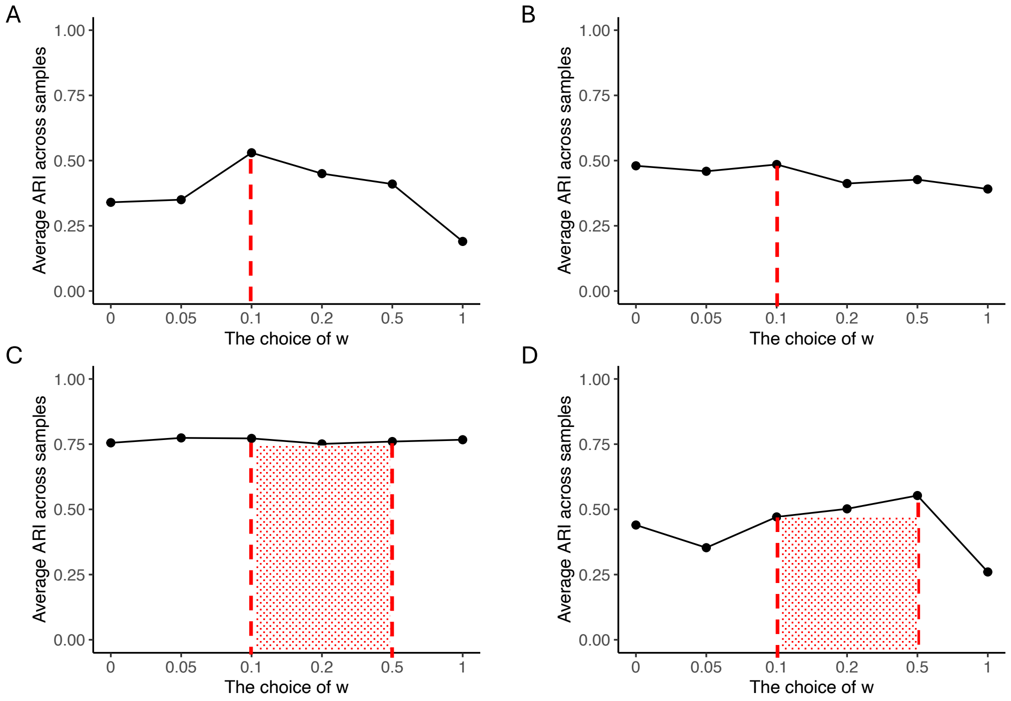


Figure S4. Sensitivity analysis of $w$. The average ARI across samples achieved by BayeSMART under different image profile weights $w$ for (A) HER2-positive breast cancer ST dataset, (B) human DLPFC 10x Visium dataset, (C) mouse mPFC STARmap dataset, and (D) mouse hypothalamus MERFISH dataset.

#### S9. More Results on the HER2-positive Breast Cancer Dataset

When classifying the spatial domains as cancer or non-cancer regions, the spatial domain identification task can also be treated as a binary classification problem. Performance can be evaluated using the area under the curve (AUC), F1-score, and accuracy (ACC). Table S4 shows the results of BayeSMART and BASS on the HER2-positive breast cancer dataset with these three metrics.

| AUC | | | | | |
|---|---|---|---|---|---|
| | A1 | B1 | C1 | H1 | Average |
| BayeSMART | 0.566 | 0.949 | 0.897 | 0.914 | **0.832** |
| BASS | 0.500 | 0.979 | 0.577 | 0.649 | 0.676 |
| F1-score | | | | | |
| | A1 | B1 | C1 | H1 | Average |
| BayeSMART | 0.949 | 0.857 | 0.970 | 0.858 | **0.908** |
| BASS | 0.942 | 0.949 | 0.885 | 0.484 | 0.815 |
| ACC | | | | | |
| | A1 | B1 | C1 | H1 | Average |
| BayeSMART | 0.904 | 0.922 | 0.952 | 0.887 | **0.916** |
| BASS | 0.890 | 0.975 | 0.801 | 0.742 | 0.852 |

Table S4. The result of BayeSMART and BASS on the HER2-positive breast cancer dataset with three binary classification metrics.

**S10. Result of BayeSMART on All 12 DLPFC Slides Together**

We also ran BayeSMART on all 12 slides from three donors together. The average ARI on 12 slides is 0.410, and the details of the results are shown in Figure S5A. The result shows the robustness of BayeSMART on multiple samples from different individuals. However, since our model requires an image profile as input, and there is currently no deep-learning-based model that performs well on H&E-stained images of the DLPFC, we opted to use the reference-free method, STDeconvolve. Unfortunately, STDeconvolve is not designed for multi-sample analysis, and analyzing each sample individually, even with the same number of anonymous cells set for each, leads to different representations across samples. To ensure consistency, where each anonymous cell represents the same entity across samples, they must be combined and analyzed together using STDeconvolve. However, because STDeconvolve is not intended for multi-sample use, applying it to multiple donors can introduce batch effects and potentially lead to unsatisfactory results. The UMAP plot on the left in Figure S5B clearly demonstrates a significant batch effect between different donors, while the right UMAP plot shows that the batch effect between slides within donor 1 is much smaller compared to that between donors. Therefore, we recommend using STDeconvolve exclusively on samples from the same individual rather than across multiple individuals.

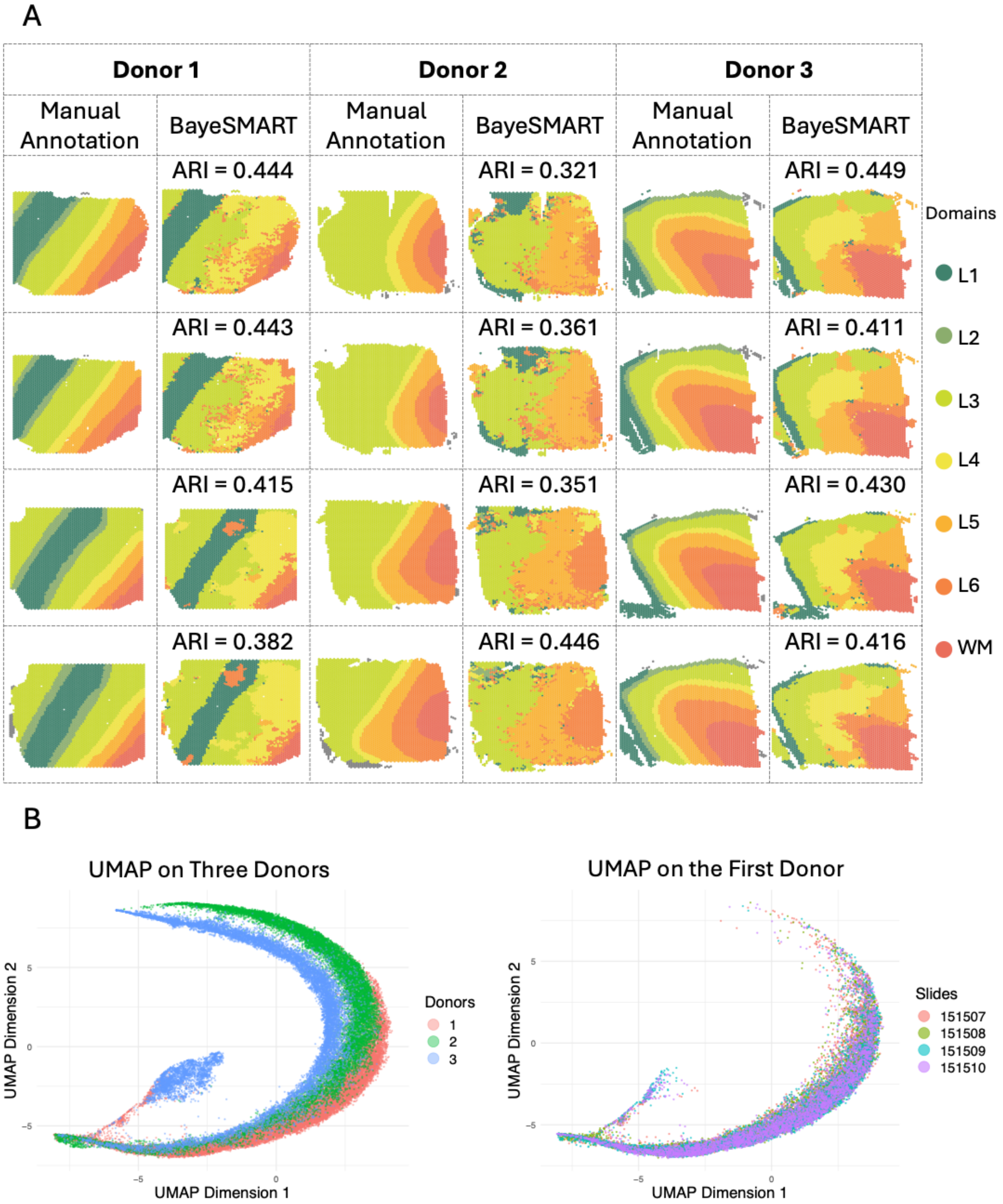


Figure S5. BayeSMART results on all 12 DLPFC slides. (A) Spatial domain identification results for each of the 12 slides from three donors. (B) UMAP of the gene expression count dataset from the three donors (left) and one the first donor (right).

**S11. Implementation Details of the Competing Methods**

**BASS:** BASS is a Bayesian based method that provides the spatial domain identification and cell type clustering analysis on multi-sample sinlge-cell SRT data. The details of the implementation are provided in the tutorial https://zhengli09.github.io/BASS-Analysis/. Hence, we followed their tutorials and obtained the results in the four real applications.

**STAGATE:** STAGATE is a region segmentation approach based on graph attention autoencoder that uses the gene expressions and spot coordinates. The Python code of STAGATE is publicly available on GitHub https://github.com/zhanglabtools/STAGATE. Note that the normalizing and the logarithmic transformation for the SRT raw count matrix is conducted before employing the main function “train STAGATE”. The arguments in the main function include the preprocessed gene expression data with coordinate information (adata), the weight of cell type-aware spatial neighbor network (alpha, its default is zero), and number of total epochs in training (n_epochs = 500). STAGATE employs “mclust R” to conduct clustering using R package mclust.

**iIMPACT:** iIMPACT is a Bayesian spatial domain identification method that integrates the gene expression profile, spatial information, and detailed morphological information extracted from the artificial intelligence (AI)-reconstructed histology image from a single-sample SRT experiment. An open-source implementation of the iIMPACT algorithm in R/C++ is available at https://github.com/Xijiang1997/iIMPACT. For the image profile of HER2-positive and DLPFC datasets, we applied the same preprocessing steps as for BayeSMART. For the STARmap and MERFISH dataset, we follow the default setting of the method provided by iIMPACT.

**BayesSpace:** BayesSpace is a Bayesian spatial domain identification method that only leverages ST data. The R code of BayesSpace is publicly available on https://github.com/edward130603/BayesSpace. The arguments in the main function spatialCluster() include preprocessed gene expression matrix with spot coordinates stored in SingleCellExperiment object (sce), predetermined region number (q), number of MCMC iterations (nrep), number of iterations in the burn-in period (burn.in), spatial transcriptomic platform (platform), and method for initialization (init.method). In the real applications, we let the initialization method to be Kmeans and set the predetermined region number as the number of domains in the annotation. The other arguments are fixed at the default values.

**DRSC:** DRSC is a frequentist approach for spatial domain identification of SRT data. The main function is available in the R package DR.SC. Firstly, we use the function find_neighbors2() to construct an adjacent neighborhood matrix with different platforms (ST or Visium). In the main function DR.SC_fit(), the number of clusters K is the set to be the number of domains in the annotation.

**Leiden:** Leiden is a graph-based community detection method, integrated into function FindClusters() in the R package Seurat. With grid search method, the resolution parameter was selected from {0.1, 0.2, …, 1.5} to ensure the number of clusters equal to the number of domains in the manual annotation of the real SRT datasets. The clustering result was obtained under the chosen resolution parameter.

**Louvain:** Louvain is an industry-standard method widely utilized in single-cell RNAseq data analysis. It is a graph-based community detection method, integrated into the R package Seurat. According to the grid search method, the resolution parameter was selected from {0.1, 0.2, . . ., 1.5} to ensure the number of clusters equal to true values in the manual annotation. The clustering result was obtained under the chosen resolution parameter.

**SCMEB:** SCMEB is a frequentist approach for spatial domain identification of SRT data. The main function is available at R package SC.MEB. Firstly, we use the function find_neighbors2() to construct an adjacent neighborhood matrix with different platforms (ST or Visium). In the main function SC.MEB(), argument K is the set of candidates for the number of clusters. In the real applications, K is the number of clusters and is set to be the same as in the manual annotation. Other arguments are default settings.

**SpaGCN:** SpaGCN is a graph-convolutional-network-based method integrating gene expression, spatial location, and histology to identify spatial domains and spatially variable genes. The Python code of SpaGCN is publicly available on GitHub https://github.com/jianhuupenn/SpaGCN. The tutorial in https://github.com/jianhuupenn/SpaGCN/blob/master/tutorial/tutorial.md provides the details of the implementation where the prespecified number of clusters is set as the number of clusters in the manual annotation.

**S12. Quality Control on HER2-positive Breast Cancer Dataset**

Since BayeSMART relies on image profiles generated by existing deep learning models for nuclei segmentation and classification on H&E-stained tissue slide images, the accuracy of cell type detection heavily depends on the quality of these images. For the HER2-positive dataset, we have eight samples with manual annotations from eight different individuals. However, in datasets like this, the image quality may vary among samples and may not always meet the required standards, especially since most articles providing such data are not primarily intended for training image models. Upon careful assessment, we found that the staining in images E and F was relatively darker than in the other samples, making it difficult to accurately identify cells. Due to these quality concerns, we excluded images E and F from our analysis. Images D and G have similar issues, and their manual annotations are of poorer quality compared to the other four images, as suggested by our collaborating pathologist. To ensure an accurate ground truth for calculating ARIs, we removed these samples from the study. Figure S10 illustrates the comparison between high-quality and low-quality images.

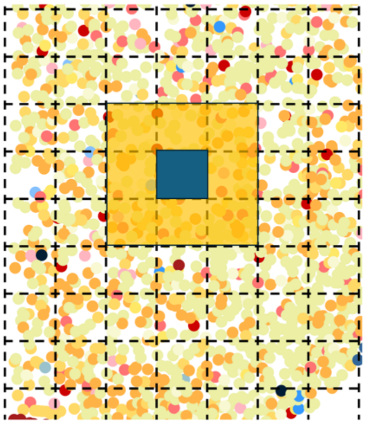

Figure S6. The neighbors of a spot within the same section for the single-cell resolution SRT data. The spots in the yellow shade are neighbors of the blue spot.

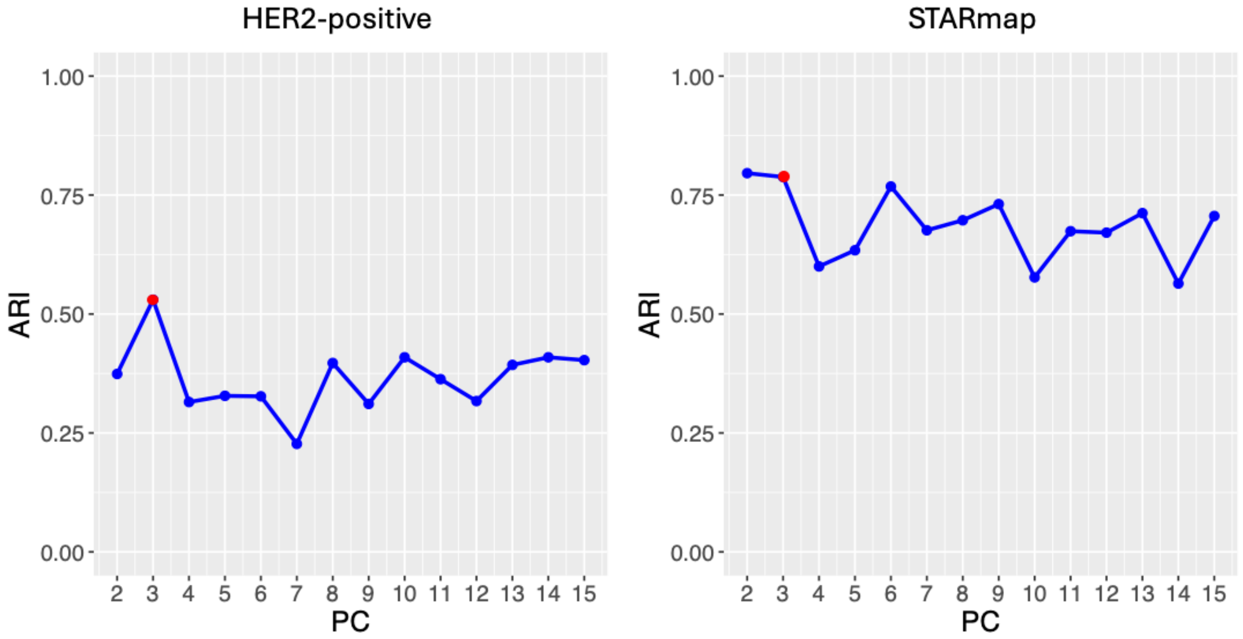


Figure S7. Sensitivity analysis on the choices of the number of principal components (PCs). The ARIs achieved by BayeSMART are shown for different numbers of PCs in the PCA for the HER2-positive and STARmap datasets. Notably, when the number of PCs is set to three, both spot-level SRT data (HER2-positive) and single-cell level SRT data (STARmap) achieve a high ARI.

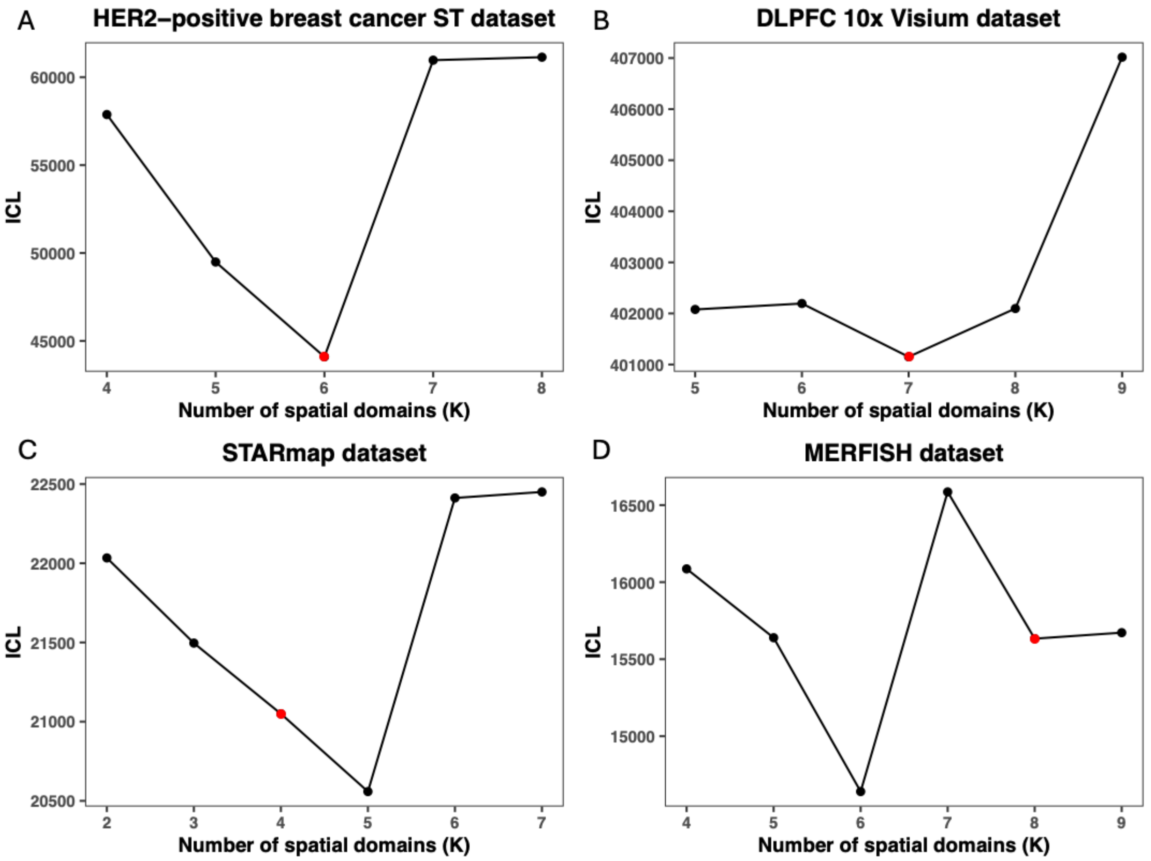


Figure S8. Integrated completed log-likelihood (ICL) plots for selecting the number of spatial domains, $K$, across different datasets: A. HER2-positive human breast cancer ST data; B. DLPFC 10x Visium data; C. mPFC STARmap data; D. mouse hypothalamus MERFISH data. The manually annotated number of spatial domains (marked in red) corresponds to the lowest ICL in the HER2-positive and DLPFC datasets, and the second lowest in the STARmap and MERFISH datasets.

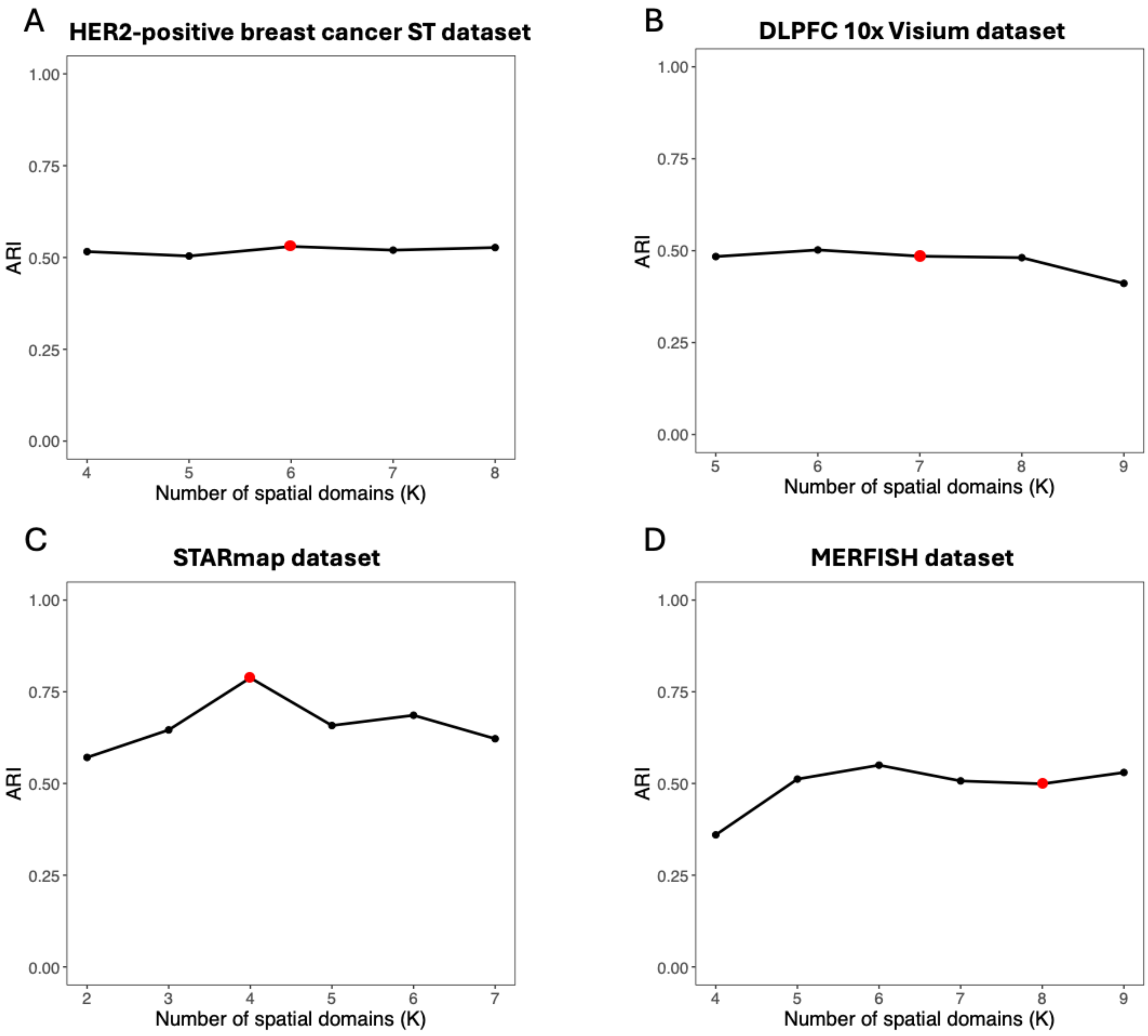


Figure S9. ARI results of BayeSMART on four datasets for different numbers of spatial domains $K$. A. HER2-positive human breast cancer ST data; B. DLPFC 10x Visium data; C. mPFC STARmap data; D. mouse hypothalamus MERFISH data. The red dots refer to the results when $K$ equals to the number of spatial domains in the manual annotation.

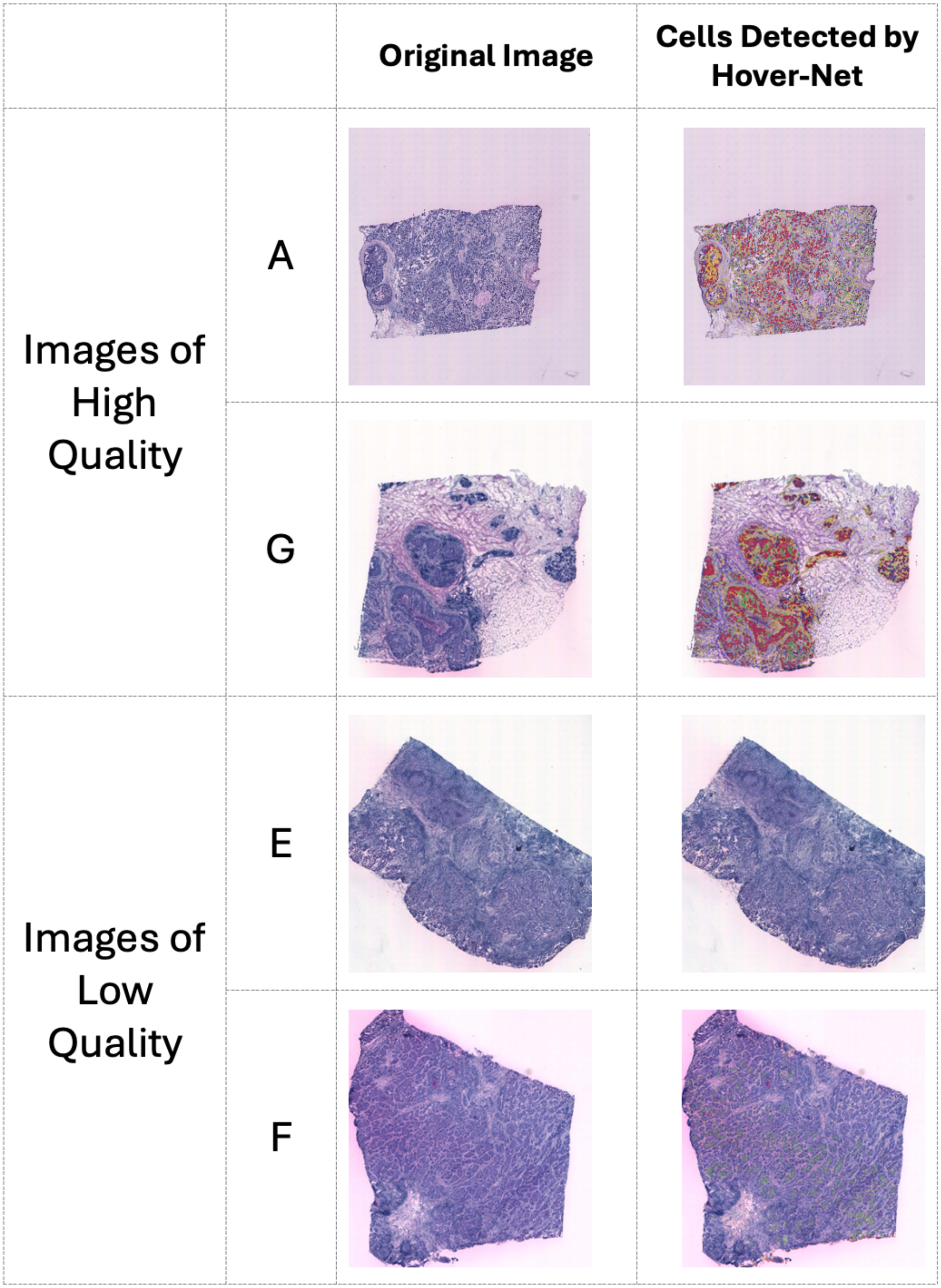


Figure S10. Comparison between high-quality and low-quality images. Cells can be detected by Hover-Net in the H&E-stained images of patients A and G; however, for images E and F, the darker tone makes it more difficult for Hover-Net to detect cells.
